# Polygenic adaptation to an environmental shift: temporal dynamics of variation under Gaussian stabilizing selection and additive effects on a single trait

**DOI:** 10.1101/505750

**Authors:** Kevin R. Thornton

## Abstract

Predictions about the effect of natural selection on patterns of linked neutral variation are largely based on models involving the rapid fixation of unconditionally beneficial mutations. However, when phenotypes adapt to a new optimum trait value, the strength of selection on individual mutations decreases as the population adapts. Here, I use explicit forward simulations of a single trait with additive-effect mutations adapting to an optimum shift. Detectable “hitch-hiking” patterns are only apparent if i. the optimum shifts are large with respect to equilibrium variation for the trait, ii. mutation rates to large-effect mutations are low, and iii., large-effect mutations rapidly increase in frequency and eventually reach fixation, which typically occurs after the population reaches the new optimum. For the parameters simulated here, partial sweeps do not appreciably affect patterns of linked variation, even when the mutations are strongly selected. The contribution of new mutations versus standing variation to fixation depends on the mutation rate affecting trait values. Given the fixation of a strongly-selected variant, patterns of hitch-hiking are similar on average for the two classes of sweeps because sweeps from standing variation involving large-effect mutations are rare when the optimum shifts. The distribution of effect sizes of new mutations has little effect on the time to reach the new optimum, but reducing the mutational variance increases the magnitude of hitch-hiking patterns. In general, populations reach the new optimum prior to the completion of any sweeps, and the times to fixation are longer for this model than for standard models of directional selection. The long fixation times are due to a combination of declining selection pressures during adaptation and the possibility of interference among weakly selected sites for traits with high mutation rates.

## Introduction

Empirical population genetics seeks to understand the evolutionary histories of natural populations by analyzing genome-wide patterns of polymorphism. The interpretation of observed patterns relies heavily on mathematical models, accompanied by various simulation methods, which make concrete predictions about the effect of evolutionary forces (natural selection, demographic events, etc.) on patterns of variation.

The models of natural selection used to interpret data come primarily from what we may call “standard population genetics” models. In these models, mutations have a direct effect on fitness (a “selection coefficient”). The fitness effects of mutations are most often assumed to be constant over time. For example, background selection is a model of unconditionally deleterious mutations resulting in strong purifying selection (Charlesworth *et al.*, 1993, 1995; Hudson and Kaplan, 1995; CvijoviĆ *et al.*, 2018). The model of a selective sweep from a new mutation similarly posits that the variant is unconditionally beneficial with a constant effect on fitness over time (Maynard-Smith and Haigh, 1974; Kaplan *et al.*, 1989; Braverman *et al.*, 1995; Durrett and Schweinsberg, 2004), and a similar assumption is made in models of selection from standing genetic variation (Hermisson and Pennings, 2005; Berg and Coop, 2015).

The effect of natural selection on linked neutral variation has been extensively studied for the case of directional selection on mutations with direct effects on fitness (*e.g.* Stephan *et al.*, 1992; Wiehe and Stephan, 1993; Kaplan *et al.*, 1989). This framework leads to a natural simulation scheme using the structured coalescent (Kaplan *et al.*, 1988) which has been widely used to study the power of various approaches to detect recent sweeps from new mutations (Fay and Wu, 2000; Kim and Nielsen, 2004), from standing variation (Prze-worski *et al.*, 2005; Innan and Kim, 2004; Hermisson and Pennings, 2005), from new mutations occurring at a fixed rate in the genome (Braverman *et al.*, 1995; Przeworski, 2002), or for testing methods to distinguish between various models of adaptation (Garud *et al.*, 2015; Schrider and Kern, 2016).

The model of Gaussian stabilizing selection around an optimal trait value differs from the standard model in that mutations affect fitness *indirectly* via their effects on trait values. For the additive model of gene action considered here, and considering a single segregating mutation affecting the trait, the mode of selection is under-or over-dominant in a frequency-dependent manner (Robertson, 1956; Kimura, 1981). This model has been extended to multiple mutations in linkage equilibrium by several authors (Barton, 1986; de Vladar and Barton, 2014; Jain and Stephan, 2015, 2017b).

The equilibrium conditions of models of Gaussian stabilizing selection on traits have been studied extensively (BÜrger, 2000, chapters 4 and 5). In general, the dynamics are quite complicated, with many possible equilibria existing for the case of many biallelic loci with equal effect sizes and no linkage disequilibrium (Barton, 1986). It is also common to assume that the forwards and backwards mutation rates per locus are equal (Barton, 1986; de Vladar and Barton, 2014; Jain and Stephan, 2015, 2017b). Under these assumptions, and assuming distributions of mutational effects symmetric around zero, large-effect variants (*e.g.* those with effect sizes greater than 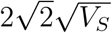, where *V*_*S*_ is the variance of the Gaussian fitness function) will be near the boundaries while small-effect variants will be at frequencies near one-half (de Vladar and Barton, 2014; Jain and Stephan, 2017b). Here, “large” and “small” effect is with respect to the effect of a variant on the genetic load of a population (de Vladar and Barton, 2014).

While the fitness effects of individual mutations on trait values affects their fixation probabilities, change in the mean phenotype of a population depends on the additive genetic variance (Robertson, 1960). When most mutational effects are small and additive, fixations require on the order of the population size in generations because phenotypic change proceeds via the fixation of small effect mutation primarily by genetic drift (Robertson, 1960). Recent theoretical work has attempted to clarify when sweeps should happen and when adaptation should proceed primarily via subtle allele frequency shifts. Chevin and Hospital (2008) considered the case of a single mutation with a large effect on fitness in a highly polygenic background evolving according to an infinitesimal model. The authors found that sweeps stall at intermediate frequencies because frequency shifts in the polygenic background contribute to adaptation. Under models of linkage equilibrium, additive mutational effects, and equal rates of forward and back mutation at a biallelic locus (Barton, 1986; de Vladar and Barton, 2014), polygenic traits adapt quickly to a sudden shift in the optimum via directional selection (Jain and Stephan, 2017b). In an infinitely large population, mutations that are rare at the time of the optimum shift may fix if their effect sizes are not overly large relative to the magnitude of the shift. The number of large-effect sweeps during adaptation depends on the magnitude of the shift and the average effect size of segregating variants (Jain and Stephan, 2017b). After the directional phase, selection becomes disruptive and mutations affecting fitness are fixed or lost in order to reduce the genetic load of the population.

Under a model of a trait with a small number of phenotypic classes, Höllinger *et al.* (2019) describe the dynamics of mutations following an optimum shift for traits with low mutation rates and for highly polygenic traits. The key parameter in their model is Θ = 4*Nµ*, where *µ* is the mutation rate relevant to the trait. When Θ ≲ 1, adaptation primarily occurs via complete sweeps. At intermediate values (Θ ≈ 10), partial and complete sweeps occur by the time the population has adapted. When Θ ≈ 100, adaptation (defined as when mean fitness has recovered following the optimum shift) proceeds via frequency shifts at many loci.

While the work described above identifies the conditions where sweeps are expected, we do not have a picture of the dynamics of linked selection during adaptation to an optimum shift. In large part, the difficulty of analyzing models of continuous phenotypes with partial linkage among sites has been an impediment to a theoretical description of the process. In general, the standard model of a single trait with additive effect mutations and Gaussian stabilizing selection assumes linkage equilibrium (or quasi-linkage equilibrium) (Turelli, 1984; Barton, 1986; de Vladar and Barton, 2014; Jain and Stephan, 2015, 2017b). Höllinger *et al.* (2019) were able to accommodate partial linkage by simplifying how mutations affect phenotype and focusing on the dynamics up until a particular mean trait value was first reached. In their simplest model, an individual is either mutant or non-mutant, and therefore there are only two phenotypes possible.

Here, I use explicit forward-time simulations to describe the average dynamics of linked selection during the adaptation of a single trait under “real” stabilizing selection (Johnson and Barton, 2005) as it adapts to a single, sudden, shift in the optimum trait value. These simulations accommodate genetic drift and partial linkage, and are also able to track the dynamics of neutral variants over time. By restricting mutations affecting the trait to specific “loci” (within which linkage is still relatively loose), and allowing neutral mutations to occur over much larger genomic intervals containing the loci, I describe the physical distances over which hitch-hiking during polygenic adaptation leaves detectable signatures. The simulations conducted here are therefore analogous to those used to study the spatial dynamics of linked selection via the structured coalescent (Kaplan *et al.*, 1988; Braverman *et al.*, 1995; Kim and Stephan, 2002; Przeworski, 2002). The key conceptual difference is that the model of adaptation is changed from constant directional selection to the sudden optimum shift models involving a continuous trait considered in de Vladar and Barton (2014); Jain and Stephan (2015, 2017b). I also investigate the effect of the recombination rate on the time to adaptation and the fixation time of beneficial mutations with respect to the mean time required to adapt to the new optimum.

## Methods

### Modeling stabilizing selection

I model a single trait under “real” stabilizing selection (Johnson and Barton, 2005). Mutations affecting trait values arise at rate *µ* per haploid genome per generation according to an infinitely many sites scheme (Kimura, 1969). For the majority of results, the effect sizes of new mutations on trait values, *γ*, are drawn from a Gaussian distribution with mean zero and standard deviation *σ*_*γ*_. Mutations have additive effects on trait value and therefore an individual’s genetic value, *z*, is the sum of all effect sizes in that individual.

Here, I use the term “locus” to refer to a continuous genomic region within which mutation and recombination events occur uniformly. Within a locus, mutations occur at positions according to a uniform continuous distribution according to an infinitely many sites scheme. Thus, each mutation results in a biallelic variant and, in the case of trait-affecting mutations, the derived allele affects trait values. What I refer to here as mutations are typically referred to as loci in much of the theoretical literature (Turelli, 1984; Barton, 1986; Robertson, 1956, 1960; de Vladar and Barton, 2014; Jain and Stephan, 2015, 2017b).

Traits are under Gaussian stabilizing selection, such that fitness, *w*, is 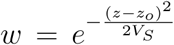, where *z*_*o*_ is the optimal trait value and *V*_*S*_ is the sum of the variance in fitness plus the environmental variance in phenotype (BÜrger, 2000, p. 160). Figure 1 shows a schematic of the model. For all simulations performed here, I used *V*_*S*_ = 1.

**Figure 1:**
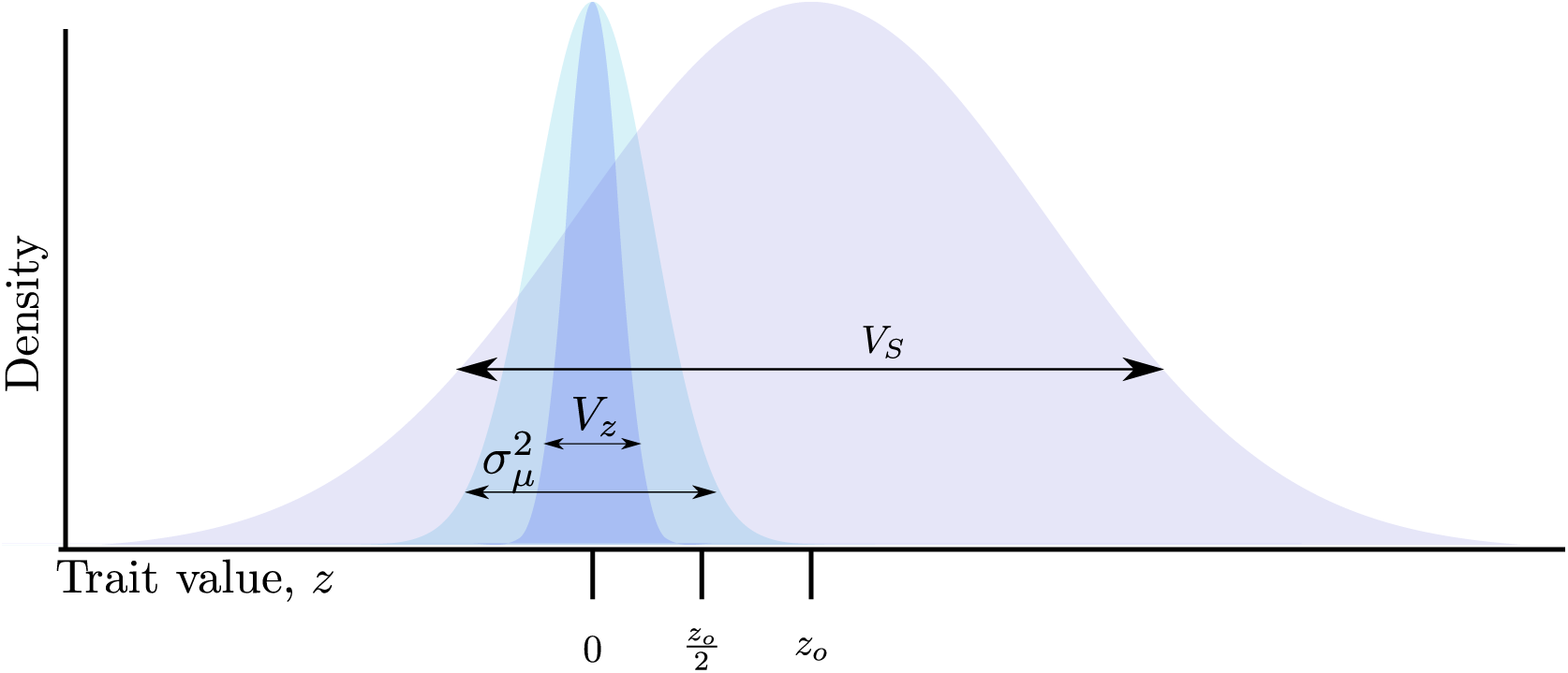
Schematic of the model. A Wright-Fisher population evolves to equilibrium around an optimum trait under Gaussian stabilizing selection with mean zero, where the parameter *V*_*S*_ represents the intensity of selection against extreme trait values 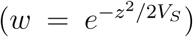. At equilibrium, the mean trait value is 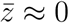 and the genetic variance, *V*_*G*_ equals the phenotypic variance *V*_*z*_. Mutations arise at a constant rate with effect sizes, *γ*, drawn from a Gaussian distribution with mean zero and variance 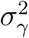. The optimum then shifts to *z*_*o*_ *>* 0, such that 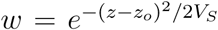. During adaptation, 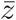 approaches *z*_*o*_ due to allele frequency change and new mutations. At any point during adaptation, mutation with effect sizes 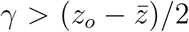 will overshoot the optimum if they reach high frequency or fix.

I model an environmental challenge as sudden “optimum shift”, where the optimum trait value changes from *z*_*o*_ = 0 to *z*_*o*_ *>* 0.

It is important to note that I am considering all of the heritable variation for the trait to be modeled in the genomic regions that are explicitly simulated. Thus, the approach is similar in spirit to that of de Vladar and Barton (2014), but with partial linkage. An alternative would be to allow for a genetic background that also evolves, for which we are not tracking mutation fates. Chevin and Hospital (2008) used the latter approach to mathematically model the dynamics of large-effect mutations in an infinitesimal background and Stetter *et al.* (2018) used a simple version of this method to simulate the dynamics of quantitative traits evolving under truncation selection.

### Forward simulation schemes

I ran all simulations using two different Python packages (see Software availability below) based on the C++ library fwdpp (Thornton, 2014). For a given diploid population size, *N*, I simulated for 10*N* generations with *z*_*o*_ = 0, at which point the optimum shifts and evolution continued for another 10*N* generations.

#### Simulating large genomic regions with only selected variants

To study the dynamics of mutations affecting trait values over time, I evolved populations of size *N* = 5, 000 diploids where mutations affecting trait values occur uniformly (at rate *µ*) in a continuous genomic interval in which recombination breakpoints arise according to a uniform Poisson process with a mean of 0.5 recombination breakpoints per diploid. The mutation rates used were 2.5 × 10^*-*4^, 10^−3^ and 5 × 10^−3^, which is the total mutation rate per haploid genome. The total mutation rate per diploid, *U*, is 2*µ*. These mutation rates correspond to Θ = 4*Nµ* values of 5, 20, and 100, respectively, meaning we expect sweeps to high frequency, mixes of partial and complete sweeps, and adaptation primarily by allele frequency changes, respectively, as the population approaches the new optimum (HÖllinger *et al.*, 2019). The three post-shift optima used were *z*_*o*_ = 0.1, 0.5, and 1. For all combinations of *µ* and *z*_*o*_, *V*_*S*_ = 1, and *σ*_*γ*_ = 0.25. At mutation-selection equilibrium, these parameters result in an equilibrium genetic variance given by the “House of Cards” approximation, which is ≈ 4*µ* for the definition of mutation rate and the *V*_*S*_ used here, and ignoring the contribution of genetic drift (Turelli, 1984). With drift, the expected *V*_*G*_ differs from the deterministic approximation by a factor of 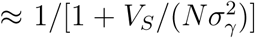 (BÜrger (2000), p. 270, equation 2.8), which is ≈ 1 for the parameters used here. For the low *µ*, low *V*_*S*_ used here, the expected genetic variance is therefore small and new mutations are more likely to have large effects relative to standing variation.

For the mutation rates and *σ*_*γ*_ defined above, the mutational variances of the trait are 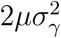, or 3.25 × 10^−5^, 1.25 × 10^−4^, or 6.25 × 10^−4^, respectively, for each mutation rate. In practice, mutational variances are often estimated with respect to the environmental variants, which poses a small issue in relating the parameters to available estimates. Here, we simulate all traits with *V*_*S*_ = 1 and do not explicitly model random effects on trait values. If instead, we were to simulate a trait with environmental variance equal to the expected genetic variance and hold *V*_*S*_ = 1, the heritability of the trait would be one-half and the evolutionary dynamics would be unaffected because the contribution of the environmental variance to *V*_*S*_ would be small (because the genetic variances simulated here are small with respect to the total *V*_*S*_). Assuming a hypothetical simulation of a trait with heritability equal to one-half, these parameters result in a ration of the mutational variance to the environmental variance of *O*(10^−2^), which is the upper limit of the ranges reported based on experimental results (Lynch (1988), Falconer and Mackay (1996), p. 349). Below, I describe simulations varying the distribution of effect sizes, thus changing the mutational variance.

For all combinations of *µ* and *z*_*o*_, various summaries of the genetic variation (*V*_*G*_, 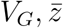, etc.) in the population were recorded every generation. In total, I ran 1,024 replicates of each parameter combination. For the first 256 replicates, the frequency trajectory of all mutations were recorded.

#### Simulating a ten locus system with neutral and selected variants

For multi-locus simulations, a locus has scaled neutral mutation rate *θ* = 4*Nµ*_*n*_ = 1, 000 and scaled recombination rate *ρ* = 4*Nr* = 1, 000, where *µ*_*n*_ is the neutral mutation rate per gamete at a locus and *r* is the mean number of recombination events per diploid at a locus. Mutation and recombination events occur uniformly along a locus and each locus is separated by 50 centi-Morgans (cM). For these simulations, I performed 256 simulation experiments per parameter combination.

Figure 2 shows how a locus is broken up into windows for analysis. Mutations affecting the trait occur in the sixth out of 11 equal-sized windows in a locus and I analyze each window separately. Thus, each window has *θ* = *ρ ≈* 90 and mutations affecting trait values are clustered in the middle of each locus (and are intermixed with neutral mutations). In these simulations, the total mutation rate affecting the trait, *µ*, is the sum over loci, and the rate per locus is equal (*µ/*10).

**Figure 2:**
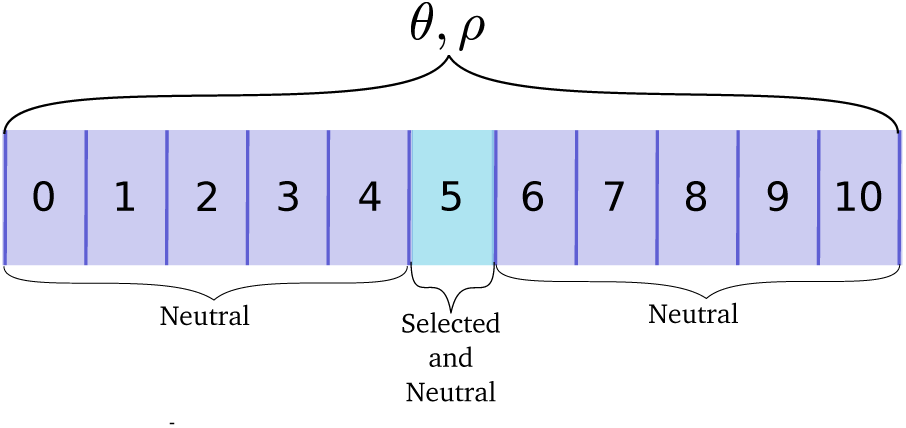
Schematic of a single locus for multi-locus simulations. The scaled neutral mutation and recombination rates, *θ* and *ρ*, respectively, are modeled as uniform processes across a locus. A locus is divided into 11 windows of equal size. Mutations affecting the trait only occur in the central window, shown in pale blue. Multiple loci are separated by 50 centiMorgans.

At each locus, mutations affecting the trait occur only in the middle window (Figure 2). Therefore, the mean number of recombination events per diploid is ≈ 0.0045 in the middle window where trait-affecting variants arise. Similarly, the mean number of new mutations per diploid at a given locus affecting the trait is *µ/*5. For the largest mutation rate used here (*µ* = 0.005), the ratio of recombination events to new mutations affecting the trait in this window is nine to one. The entire genome consists of ten such loci, for a total mutation rate of *µ* and a total *θ* = 10^4^.

For a model of a single trait under Gaussian stabilizing selection with a constant optimum, the selection coefficient 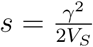 (Simons *et al.* (2018), see also Kimura and Crow (1978)). Here, *V*_*S*_ = 1, and therefore the relevant scaled strength of selection acting on a segregating variant is *Nγ*^2^. For many of the results presented here, it is helpful to treat the dynamics of strongly selected mutations separately. To do so, I define a “large-effect” variant as having *Nγ*^2^ ≥ 100, meaning that the deterministic force of selection is much stronger than that of drift. To vary the probability that a new mutation is of “large-effect”, I perform a second set of simulations, also involving ten unlinked loci, varying the distribution of effect sizes (DES) such that probability that *Nγ*^2^ ≥ 100 takes on values of 0.1, 0.5, or 0.75. For Gaussian distributions of effect sizes, the mean *γ* is zero, as above, and *σ*_*γ*_ is found by numerical optimization using `scipy` (Jones *et al.*, 2001–) to give the desired *Pr*(*Nγ*^2^ ≥ 100). I also use Gamma distributions with shape parameters equal to either one or one-half, and then find a value for the mean of the distribution using `scipy`. These shape parameters give probability density functions that are “exponential-like” in shape. For simulations with Gamma DES, I use an equal mixture of Gamma distributions with mean *γ* and *-γ* such that the DES is symmetric around a value of zero. I performed 100 simulation replicates for each parameter combination. Using the argument from above, assuming hypothetical simulations of a trait with a heritability of one-half, the Gaussian distribution and the gamma with a shape of one give a ratio of the mutational variance to the environmental variance of 2 ×10^−3^ to 3 × 10^−3^ when the proportion of new mutations with *Nγ*^2^ ≥ 100 is 0.1. These values are close to the mean of ≈ 10^−3^ reported for a variety of traits (Lynch (1988), Falconer and Mackay (1996), p. 349)

In a third set of simulations, I vary *ρ* = 4*Nr*, the recombination rate within each locus. I ran 256 replicates of these simulations using the tree sequence recording algorithm (Kelleher *et al.*, 2018) implemented in `fwdpy11` version 0.3.2. For these simulations, I recorded the entire population as nodes in the tree sequences for each of 200 generations after the optimum shift. Recording nodes from these time points allows them to be analyzed after the simulation has completed. Each replicate was simulated twice. The first run simply output metadata about mutations that reached fixation. The second run was performed with the same random number seed as the first and used the metadata from the first run to track linkage disequilibrium around fixations over time, outputting those data along with the tree sequence for the simulation.

### Genome scan statistics from multi-locus simulations

The ten-locus simulations described above were used to look at the temporal dynamics of several population-genetic summaries of a sample. Each of the ten loci consists of 11 non-overlapping windows (Figure 2) and all summary statistics are calculated on a per-window basis. I used `pylibseq` version 0.2.1 (https://github.com/molpopgen/pylibseq), which is a Python interface to `libsequence` (Thornton, 2003), to calculate all genome scan statistics. All statistics were obtained from 50 randomly chosen diploids.

#### *z*-scores for the *nS*_*L*_ statistic

Individual values of the *nS*_*L*_ statistic (Ferrer-Admetlla *et al.*, 2014) from the first and last window of each locus were binned into intervals of size 0.1 based on derived frequency. These windows were used because they are the furthest from mutations affecting trait values, and thus the least affected by linked selection. The data from all loci were combined and the mean and standard deviation of each bin were used to obtain *z*-scores for markers from the remaining windows.

#### Coalescent simulation

I used `msprime` (Kelleher *et al.*, 2016) version 0.5.0 for all coalescent simulations under neutral models and `discoal` (Kern and Schrider, 2016) version 0.1.1 for all simulations of selective sweeps. All simulation output were processed using `pylibseq` version 0.2.1.

#### Software availability

I used `fwdpy` version 0.0.4 (http://molpopgen.github.io/fwdpy) compiled against `fwdpp` version 0.5.4 (http://molpopgen.github.io/fwdpp) for single-region simulations. I used `fwdpy11` versions 0.1.4, 0.2.1,, 0.3.2, and 0.5.1 (http://molpopgen.github.io/fwdpy11) for all multi-region simulations. `fwdpy11` is also based on `fwdpp`, and includes that library’s source code for ease of installation. Both packages were developed for the current work, but only the latter will be maintained.

I used the Python package pylibseq version 0.2.1 (http://pypi.python.org/pypi/pylibseq/0.2.1), which is a Python interface to libsequence (Thornton, 2003), to calculate population-genetic summary statistics.

All of these packages are available under the terms of the GNU Public License from http://www.github.com/molpopgen. The specific software versions used here are available for Linux via Bioconda (Dale *et al.*, 2017), with the exception of `fwdpy11` 0.2.1, which must be installed from source. I have made all Python and R (R Core Team, 2016) scripts for this work available at http://github.com/molpopgen/qtrait_paper.

#### Open source tools used

Data processing and plotting relied heavily on the following open-source libraries for the `R` language (R Core Team, 2016) `dplyr` (Wickham and Grolemund, 2017), `ggplot2` (Wickham and Grolemund, 2017), `lattice` (Sarkar, 2008) as well as the following Python libraries: `pandas` (McKinney, 2017), `numpy` (Van-derPlas, 2016), `matplotlib` (Hunter, 2007; VanderPlas, 2016), and `seaborn` (http://seaborn.pydata.org). The sqlite3 library (www.sqlite.org) facilitated data exchange between Python and `R` via the `pandas` and `dplyr` libraries, respectively.

## Results

### Single region results

In this section I use simulations of a large contiguous region with mutations affecting the trait occuring uniformly throughout the region. The technical details of the simulation parameters are given in Methods. Briefly, I evolved populations for 10*N* generations to mutation-selection equilibrium around an optimum trait value of *z*_*o*_ = 0, at which point *z*_*o*_ is changed to 0.1, 0.5, or 1.0 and evolution continued for another 10*N* generations. These simulations may be viewed as similar to the numerical calculations in de Vladar and Barton (2014) and Jain and Stephan (2017b), but with loose linkage between selected variants whereas the previous studies assumed linkage equilibrium and we allow for new mutation after the optimum shift. They differ from the approach of HÖllinger *et al.* (2019) in that I simulate continuous traits and do not stop evolution once a specific mean fitness is first reached.

The mean trait value, 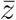, rapidly approaches the new optimum, typically reaching the new optimum within one hundred generations (Figure 3a, see also de Vladar and Barton (2014); Jain and Stephan (2017b); HÖllinger *et al.* (2019)). Prior to the optimum shift, the average genetic variance is given by 4*µV*_*S*_ (Turelli (1984), Figure 3b). Following the optimum shift, the genetic variance spikes as the population adapts (see also de Vladar and Barton, 2014; Jain and Stephan, 2017b) and then recovers to a value near 4*µV*_*S*_ within ≈ 200 generations when the mutation rate is small and taking longer to return to equilibrium when the mutation rate is higher.

**Figure 3:**
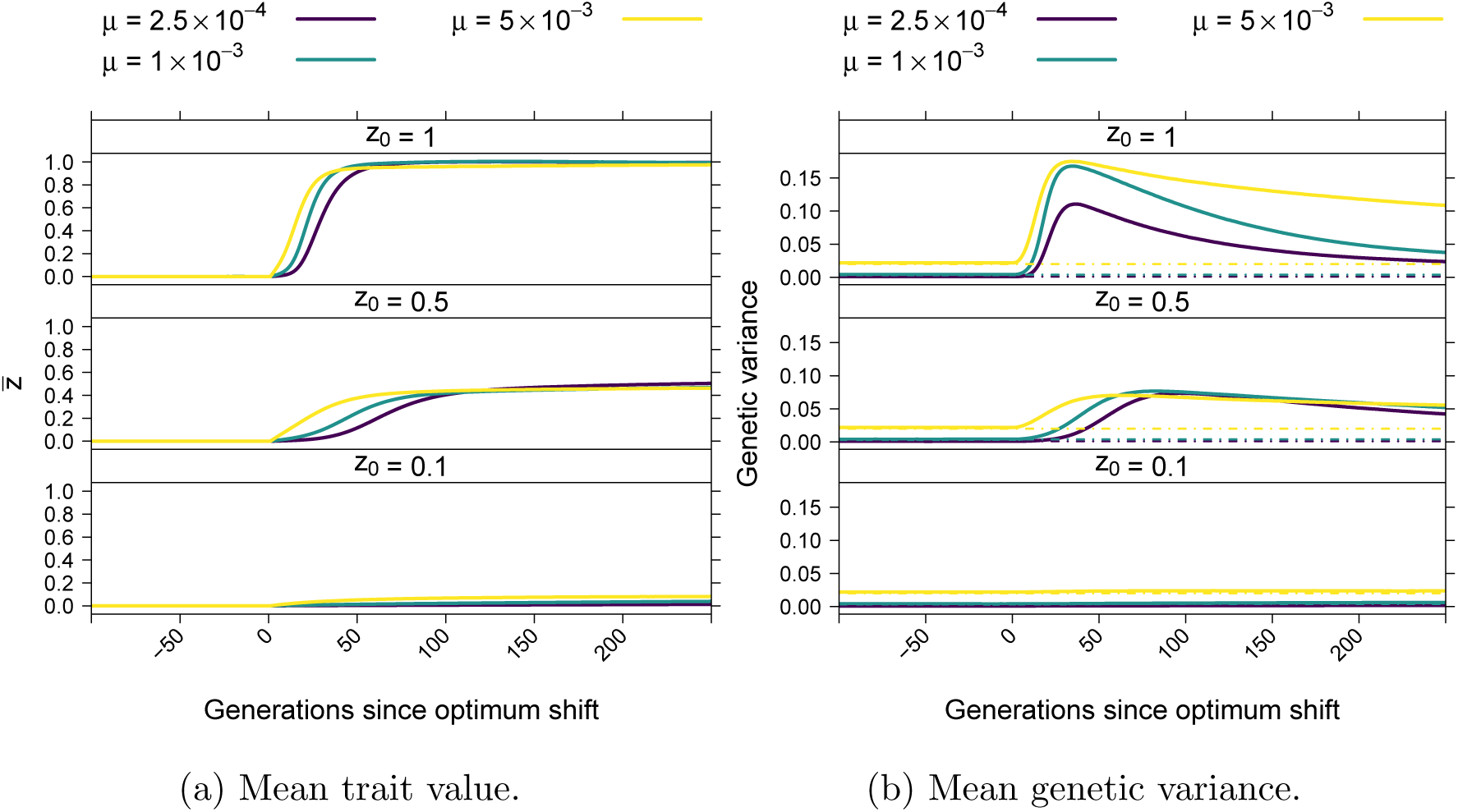
(3a) Mean trait value over time. (3b) Mean genetic variance over time. The dot-dashed lines correspond to 4*µV*_*S*_, which is the equilibrium genetic variance expected under the House-of-Cards model (Turelli, 1984).

Figure 4 shows examples of the dynamics of 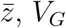, *V*_*G*_, and of mutation frequencies following the optimum shift. Each example is a single simulation replicate. The top row of plots shows that 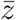 quickly reaches *z*_*o*_ for the individual replicates. The approach of 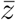to *z*_*o*_ corresponds to a substantial increase in the genetic variance similar to what is shown for the average genetic variance over time in Figure 3b. The middle row of panels in Figure 4 shows the frequency dynamics of mutations that eventually fix. Importantly, 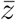 typically reaches *z*_*o*_ before the first fixation has occurred (see Figure S1 for details over a shorter time scale). The legend of of each panel in Figure 4 contains the effect sizes of variants where *Nγ*^2^ ≥ 100. The legends also contain the origin times, *o*, of these large-effect mutations, measured as generations since the optimum shift.

**Figure 4:**
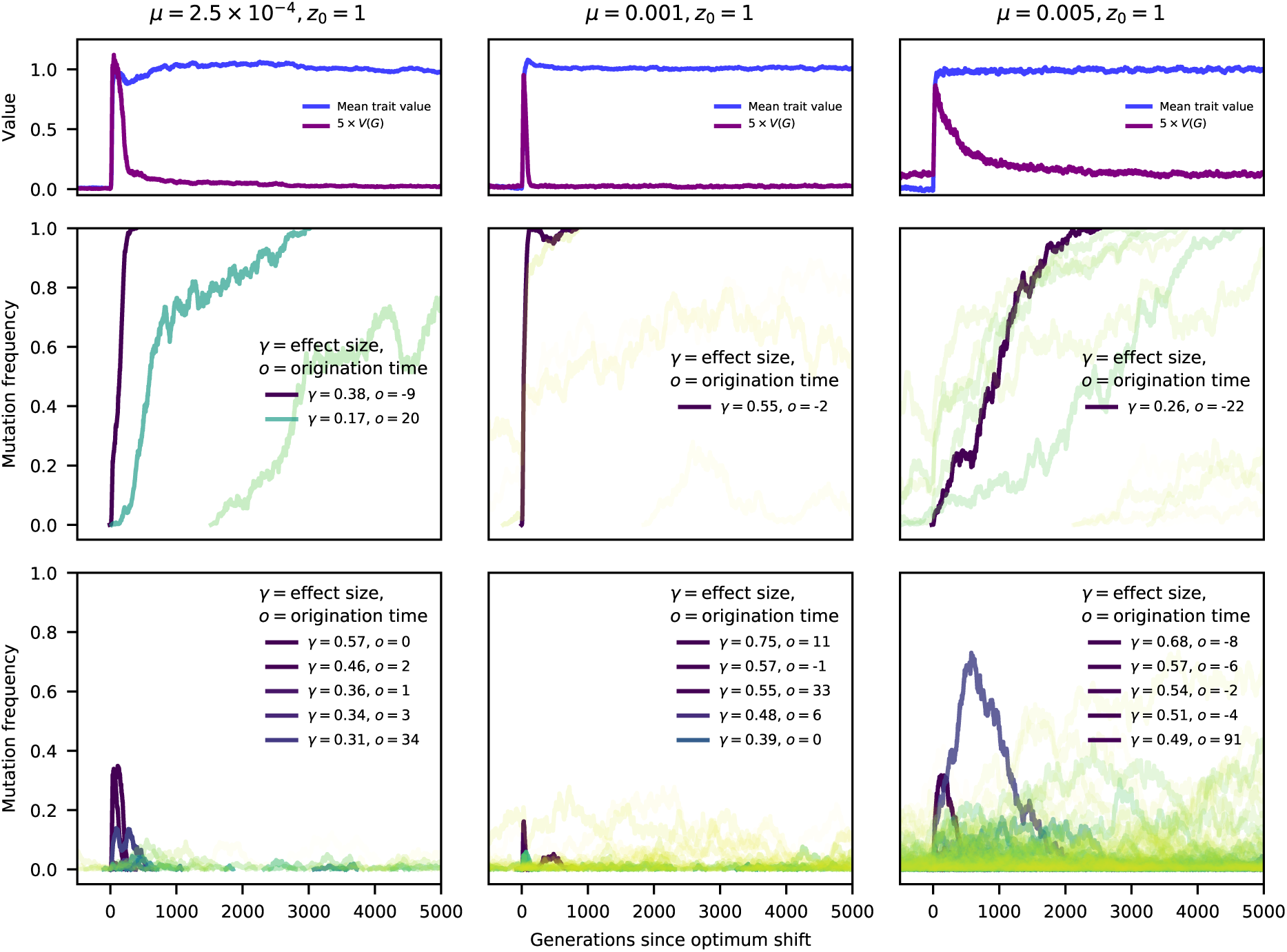
Trajectories of selected mutations. The three columns show results from a single simulation replicate for low, moderate, and high-mutation rate simulations, respectively. Parameter values are at the top of each column. The first row of plots shows the trait value and the genetic variance (multiplied by a constant for plotting purposes) over time, for up to *N* generations post optimum shift. As in Figure 3, the populations adapt quickly to the new optimum of *z*_*o*_ = 1. The middle row shows the frequency trajectories of fixations. Solid, darker (purple/blue) colors reflect larger effects on trait values, and more transparent colors in the green/yellow range reflect smaller effect sizes. Fixations with effect sizes *Nγ*^2^ ≥100 are indicated in the legend. The bottom row shows the frequency trajectories of mutations that are eventually lost. The coloration is as for the fixations, and any mutations that did not reach a frequency of one percent are excluded. A maximum of five mutations, corresponding to the five largest *|γ|* are included in the legend in the final row. Figure S1 shows the same data on a smaller time scale, showing the details on allele frequency change during the rapid adaptation to the new optimum.

For these examples, mutations with large effects on trait value fix first as predicted by Robertson (1956). In Figure 4, fixations of large effect typically have origin times close to zero, meaning that the mutation arose close to the time of the optimum shift. This observation is expected as such mutations contribute significantly to genetic load and thus their equilibrium frequencies prior to the optimum shift should be near the boundaries (de Vladar and Barton, 2014; Jain and Stephan, 2015, 2017b). Here, because of the one-way mutation model, such large-effect variants are at frequencies near zero.

The final row of plots in Figure 4 shows the dynamics of mutations that reached a frequency of at least one percent but were eventually lost from the population. Large effect mutations only exist for a relatively brief period of time after the optimum shift, after which most segregating variation reaching appreciable derived allele frequencies are of relatively small effect. An important observation in the final row of Figure 4 is that, for a short time following the optimum shift, several intermediate-frequency mutations with large effects on trait values may be segregating. Many of these variants are adaptive (*γ >* 0) but will only make short-term contributions to adaptation prior to their loss. The dynamics of these mutations recapitulate results from de Vladar and Barton (2014)–due to epistatic effects on fitness, some mutations that are initially beneficial later become deleterious and are removed. Figure S1 shows the data from Figure 4 over a shorter time scale, allowing a more detailed look at the dynamics of mutations during adaptation.

Figure 4 suggests that fixation times are rather long, on the order of *N* generations even for mutations with large *Nγ*^2^. These long fixation times are in fact typical, and large-effect mutations typically fix in *N/*2 to *N* generations (Figure S2), which is long relative to the deterministic expectation for strongly selected sweeps from new mutations (Stephan *et al.*, 1992). Large-effect mutations that reach fixation arise close to the time of the optimum shift (Figure S3) and typically show shorter fixation times (Figure S4). In general, the number of sweeps from new mutations and from standing variants are similar, although fixations of smaller-effect standing variants are more common in simulations with higher *µ* (Figure S5). In Figure S5, a sweep from a new mutation is defined as a mutation arising withing 100 generations of the optimum shift and then reaching fixation. While somewhat arbitrary, this definition is justified by the rapid mean time to adaptation (Figure 3). In this model, large-effect standing variants that fix after the optimum shift were rare at the time of the shift (Figure S6). Small-effect mutations are also typically rare at mutation-selection balance, in particular when the mutation rate is small (Figure S6).

For the parameters simulated here, and for the genetic map simulated here (Figure 2), Figures S3, S4, and S6 suggest that large-effect fixations occur from both new mutations and from standing variation, with more large-effect fixations occurring when the *µ* is smaller limited and/or the optimum shift is larger. Thus, we may predict that large-effect fixations from new mutations may show the signals of “hard sweeps”, such as an excess of high-frequency derived neutral variants (Fay and Wu, 2000; Zeng *et al.*, 2006). Given that large-effect fixations from standing variation are typically rare at the onset of directional selection (Figure S6), we may also expect them to affect linked neutral variation (Przeworski *et al.*, 2005; Berg and Coop, 2015). For the parameters simulated here, fixations from variants that are common at the time of the optimum shift have small effects on trait values (Figure S6). The fixation of such mutations are unlikely to generate the patterns of haplotype diversity associated with “soft sweeps” because such patterns require strong selection on mutations at intermediate frequencies (Garud *et al.*, 2015).

### Fitness effects of mutations during adaptation

In this section, I explore in more detail the strength of selection on individual mutations during the directional phase of selection. These dynamics are relevant to the long fixation times noted in the previous section and also to the extent to which hitch-hiking will affect patterns of linked variation, which is the topic of the next section. As the focus of the remaining sections will be on patterns of variation during adaptation, we switch from simulating a single large region to simulating ten unlinked regions. The only difference between these simulations and those described above is the genetic map and the position of mutations affecting trait values (see Methods for the technical details).

Figure 5 plots the dynamics of mutations in a ten locus system for one replicate of each of the three mutation rates used here. In each column, the gray vertical line is the time the population first reaches a mean trait value of 0.9*z*_*o*_, which corresponds to a mean fitness of at least 0.9 for each replicate. For simplicity, we will call this the time of adaptation. The top row of Figure 5a shows the frequency trajectories of mutations that eventually fixed. These replicates were chosen because each had one fixation of a strongly selected mutation with a similar effect size. As the mutation rate increases, the genetic background of these fixing variants becomes more polygenic. As a result, the initial rate of frequency change of the fixation lessens because other mutations are involved in the response to the optimum shift, some of which may contribute to adaptation but not fix in the long term. For all replicates, the fixations are at different loci (separated by at least 50cM) with one exception. For the high mutation rate case, the locus with the large-effect fixation also fixed one mutation with small *γ*.

**Figure 5:**
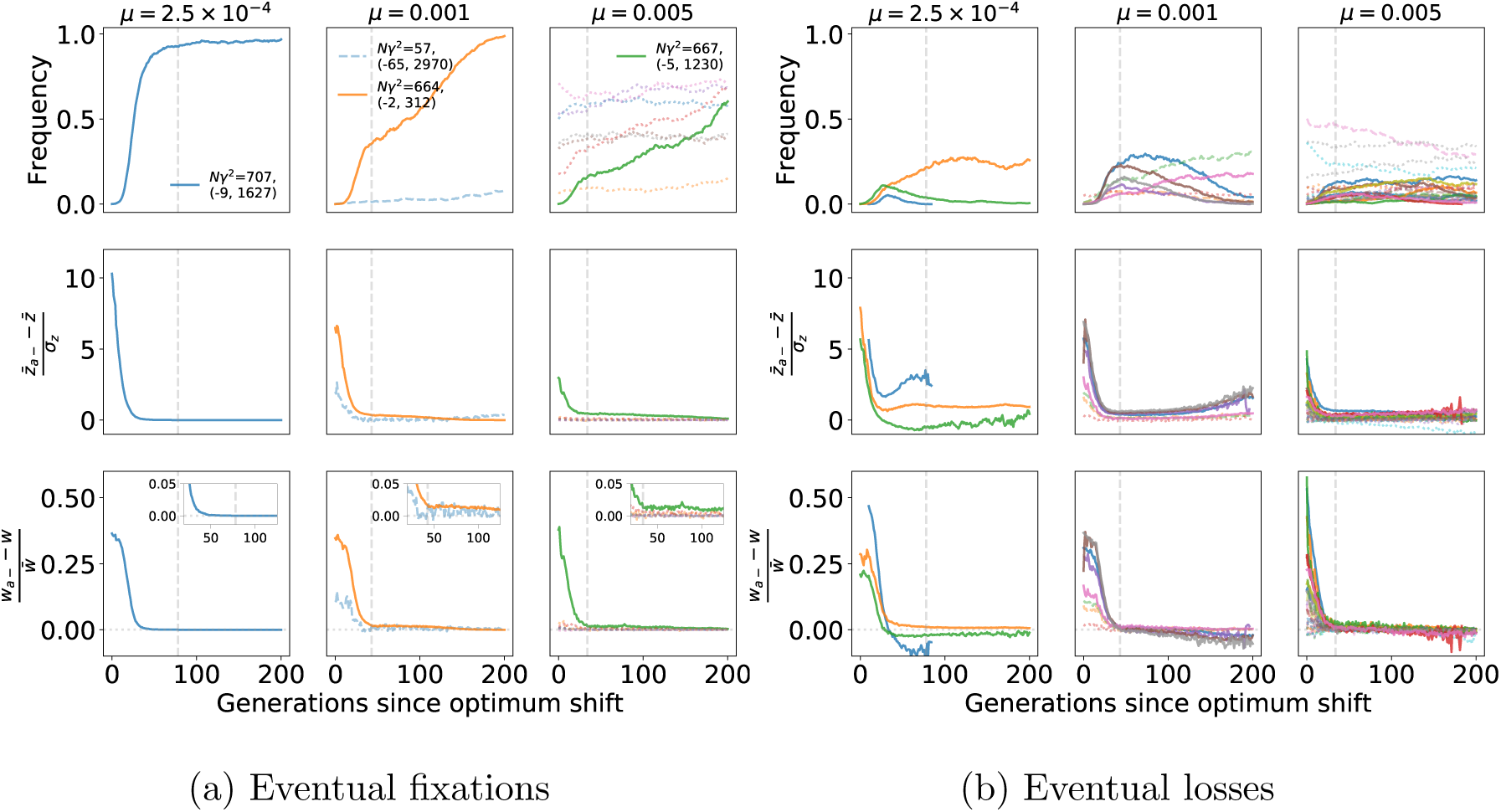
Phenotypic and fitness effects of fixations and losses. The data shown are for a single simulation replicate with *σ*_*γ*_ = 0.25, *z*_*o*_ = 1 and the mutation rate *µ* shown atop each column. The mutation rate shown is the sum over loci and individual loci mutate at equal rates (*µ/*10). In all panels, solid lines refer to *Nγ*^2^ ≥ 100, dashed lines 10 *≤ Nγ*^2^ *<* 100 and dotted lines 1 *≤ Nγ*^2^ *<* 10. The vertical line is the generation when the mean trait value first crossed 90% of the new optimal value (0.9*z*_*o*_). (a) The dynamics of fixations. The top row shows the frequency trajectory of mutations that eventually reached fixations. For mutations with *Nγ*^2^ ≥ 10, the legend shows *Nγ*^2^ and the mutation’s origin and fixation times in parentheses, scaled so that zero is the time of the optimum shift. Defining an *a–* genotype to be any genotype containing at least one copy of these “focal” mutations, the second row shows the mean deviation from the mean trait value for the focal genotypes, standardized by the phenotypic standard deviation. The final row shows the mean relative deviation in fitness for *a–* genotypes. The horizontal line in the last row is placed at the reciprocal of the population size (1*/N*). (b) The dynamics of losses. Plotting is identical to (a), but the data are filtered to only include mutations arising prior to the population first crossing 0.9*z*_*o*_ and eventually reaching a frequency of ≥ 0.05.

Figure 5b shows the frequency dynamics of mutations arising prior to adaptation that were eventually lost. As the mutation rate increases, there are more large-effect mutations increasing in frequency during adaptation. For the lowest mutation rate simulated here, two such mutations are decreasing in frequency prior to adaptation. At *µ* = 0.001, four strongly selected mutations sweep to frequencies *>* 0.10 and are later lost. For *µ* = 0.005, several large-effect mutations experience more modest increases in frequency during adaptation. From left to right, the columns of Figure 5a and 5b show that allele frequency changes are less dramatic prior to adaptation as the mutation rate increases. These results are consistent with the theoretical predictions from (HÖllinger *et al.*, 2019) that the dynamics of mutations on the time scale of adaptation are dependent on 2*NU*.

The second row in Figure 5 shows the mean deviation of a genotype with a given mutation standardized by the standard deviation in trait values (the “z”-score). The mutations that fix (Figure 5a) are all initially found in heterozygous genotypes with trait values multiple standard deviations greater than the mean. Such mutations are not necessarily the largest-effect variants present at the time of the optimum shift, which is seen for the two higher mutation rates in Figure 5. The mutations that do eventually fix were initially at higher frequency and/or associated with higher-fitness genotypes than large-effect mutations that were eventually lost.

As the population adapts, the deviation in trait value (from the population mean) for a mutation with a given effect size decreases. These *z* scores decrease because the genetic variance transiently increases following the optimum shift (Figure 3b, de Vladar and Barton (2014); Jain and Stephan (2017b)) because mutations are increasing in frequency and the variance is a function of allele frequency times the squared effect size. Mutations causing larger deviations are expected to become lost, as seen most clearly in the first column of Figure 5b–the blue and green mutations over-and under-shoot the optimum, respectively. At low mutation rates, there is a tendency to slightly over shoot on average (Figure 3) because such mutations will have larger initial increases in allele frequency than smaller-effect variants.

Finally, we can turn to the long fixation times. These are, in part, due to the decreasing strength of selection on individual mutations during the time period where directional selection occurs. The final row of Figure 5 shows the relative deviation due to genotypes carrying each mutation over time. As expected, genotypes with fitness above the mean increase in frequency, and these genotypes are associated with trait values multiple standard deviations closer to the new optimum. As the mean trait value approaches the new optimum, the relative excess fitness of these genotypes declines, approaching the reciprocal of the population size. Once the population has adapted, these mutations have small effects on phenotypic variation and their long-term dynamics are governed by underdominant selection against phenotypic variance (Robertson, 1956; Kimura, 1981). The underdominant selection means that mutations with frequencies greater than one-half will be weakly favored and are expected to fix and those with frequency less than one have will most likely be removed from the population. The small fitness differences amongst genotypes at the time of adaptation predicts that fixation times will be slow due to relatively weak selection Figure 5. Note that all of the sweeping alleles in Figure 5 are from standing variation (origin times less than zero) and are rare at the onset of directional selection (also see Figure S6).

Finally, traits with higher mutation rates have larger numbers of small-effect mutations segregating prior to adaptation (Figure 5). Once the population is adapted, the deviations from mean fitness tend to be small for most genotypes, and the large-effect mutants are not yet fixed, implying that interference (Hill and Robertson, 1966) may also increase fixation times when the mutation rate is higher. We will return to the role of interference below. The observation in Figures 4 and 5 of mutations not reaching fixation by the time the new optimum is hit consistent with previous results from other authors (HÖllinger *et al.*, 2019; Chevin and Hospital, 2008; Jain and Stephan, 2017b).

### Dynamics of linked selection in a multi-locus system

I now describe the temporal dynamics of genetic variation over time in a ten locus system. The technical details of the simulations are identical to the previous section, and are described in detail in Methods.

Figure 6 summarizes patterns of variation in the central window (Figure 2) of each locus where large-effect mutations segregate during adaptation to the new optimum. The figure is based on the data from Figure 5. The first two rows plot the frequency trajectories of eventual fixations and losses, and the next three rows summarize patterns of variation calculated from a random sample of individuals. These summaries of variation only show deviations from equilibrium values consistent with positive selection at loci where large-effect fixations occur. Further, the deviations are more pronounced when the mutation rate is smaller. The partial sweeps occurring at intermediate mutation rates (middle column of Figure 6) are not associated with strong signals of hitch-hiking, at least when the sample size is relatively small as is the case here. The time when a given statistic shows its maximum departure from equilibrium values differs for each statistic and, for the replicate with *µ* = 0.001, the maximum departure may occur ≈ 100 generations after the time to adaptation. Visually, however, one could argue that haplotype diversity tends to minimize closer to the time to adaptation than the summaries of the site frequency spectrum.

**Figure 6:**
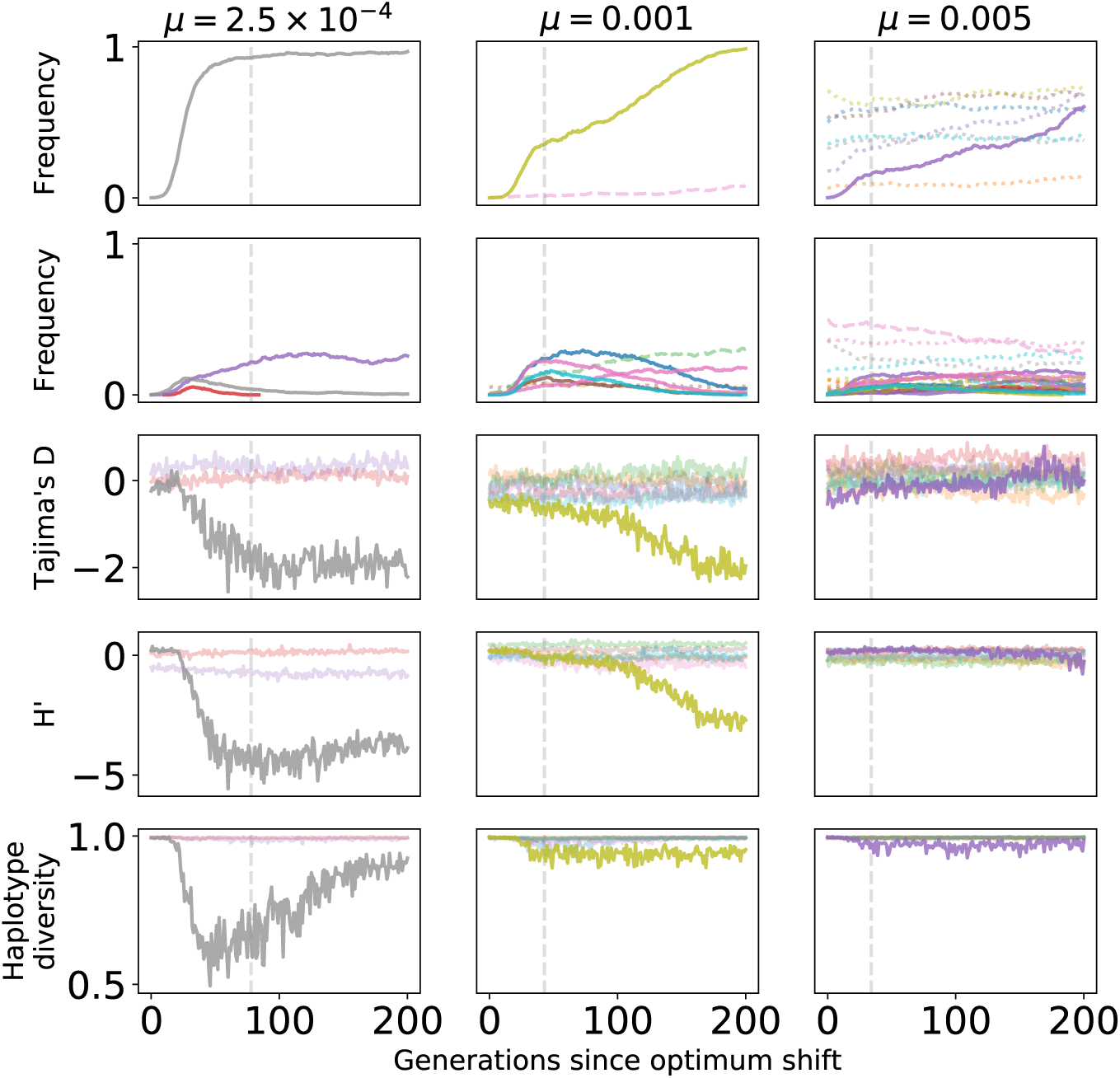
Signals of directional selection in single replicates of a ten-locus system. The data shown are based on the same simulations as in Figure 5. The first two rows show frequency trajectories for fixations and losses, respectively, with the colors indicating the locus where the mutation is found. The vertical gray line is the generation when the mean trait value first crosses 90% of the optimal trait value. The remaining rows show Tajima’s (1989) *D, H′* (Zeng *et al.*, 2006), and haplotype diversity in a random sample of 50 diploids, calculated using genotypes taken from the central “window” of a locus where causal mutations are occurring (Figure 2).

Figure 7 shows patterns of variation along each of the ten loci from an additional simulated replicate for each of the parameters shown in Figure 5 and Figure 6. Each line corresponds to a different time point in the approach to the new optimum value of *z*_*o*_ = 1, showing data for the first time the population mean trait value crosses the thresholds of 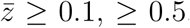 and ≥ 0.9. While the values are noisy along a genome, it is apparent that directional selection is affecting patterns of variation at linked sites in the replicates with smaller mutation rates. In the leftmost column, where *µ* = 2.5 × 10^−4^, an excess of high-frequency derived variants is seen at locus four, along with a reduction in haplotype diversity. A standing variant of large effect swept to high frequency at this locus during adaptation. In the middle column (*µ* = 10^−3^), one sees a less dramatic reduction in haplotype diversity at locus ten, where a strongly selected standing variant reached high frequency. For these two replicates, there is some evidence of reduced haplotype diversity at loci eight and five, respectively, that is not associated with any fixations. The final column, where *µ* = 5 × 10^−3^, there is no obvious temporal nor spatial pattern to variation in diversity levels, and the largest deviations from the background are not associated with the fixation of beneficial mutations.

**Figure 7:**
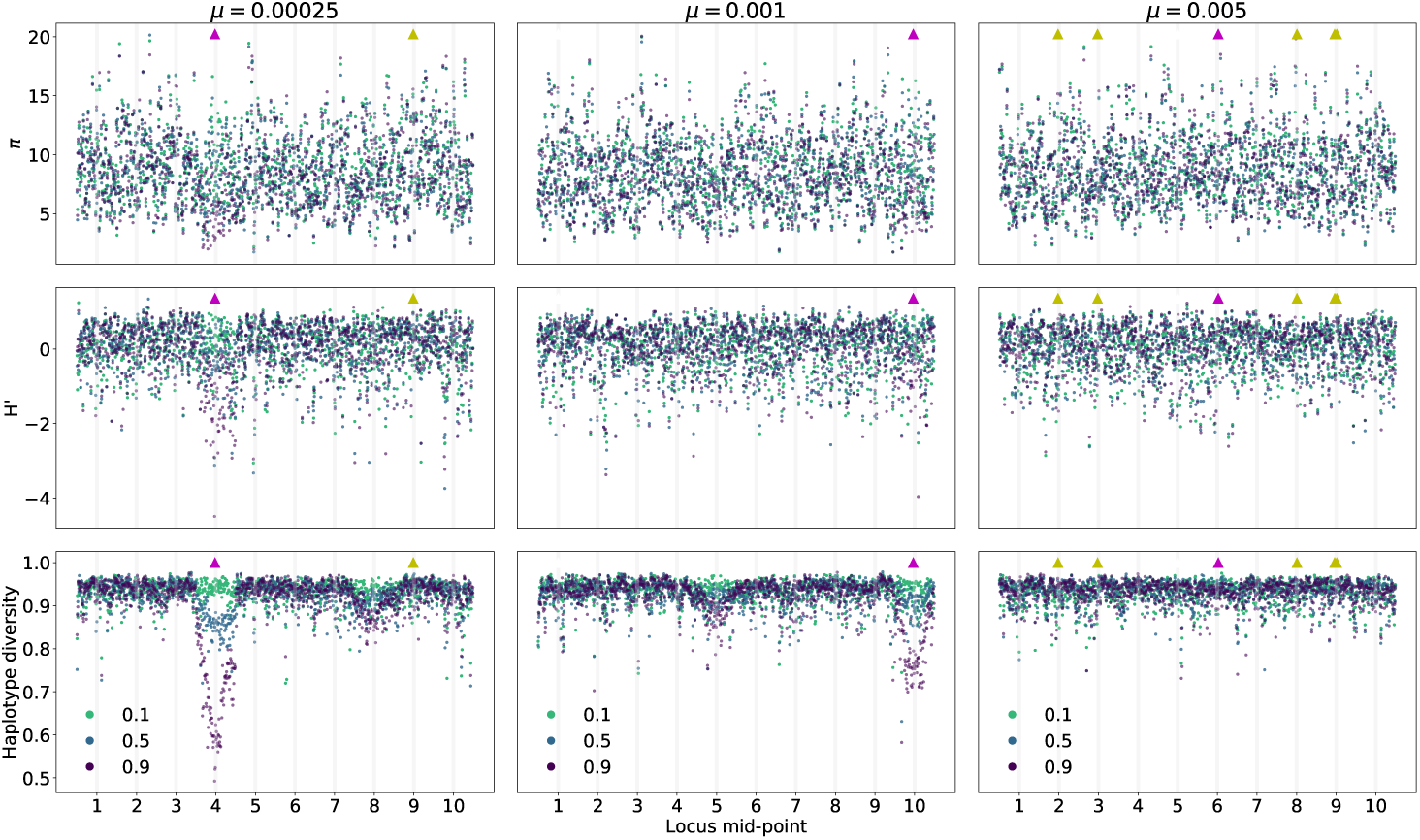
Patterns of genetic variation along genomes in a ten locus system during adaptation to an optimum value of *z*_*o*_ = 1 and *σ*_*γ*_ = 0.25. The mutation rate shown is the sum over loci and individual loci mutate at equal rates (*µ/*10). Each column corresponds to a single simulated replicate with the mutation rate given at the top. The three rows correspond to nucleotide diversity (*π*), *H′* (Zeng *et al.*, 2006), and haplotype diversity, respectively. The three point colors refer to statistics calculated from 50 randomly chosen diploids in the first generation that the population mean trait value first crossed values of at least 0.1, 0.5, or 0.9. The gray shades refer to the locations within each locus where mutations affecting trait values occur (Figure 2). The triangles along the top of each panel show where fixations occurred. Triangles pointing up are fixations from standing variation. Magenta refers to fixations with scaled effect sizes *Nγ*^2^ ≥ 100 and yellow refers to 1 *≤ Nγ*^2^ *≤* 10.

Overall, Figure 6 and Figure 7 suggest that patterns of strong hitch-hiking are more likely at loci where large-effect mutations fix. Moreover, such mutations must arise on average before the mean time to adaptation. Below, when looking at average patterns of variation over time and along genomes, we will distinguish patterns of variation where fixations meeting these conditions occur from the mean pattern expected from a randomly chosen locus.

#### The site frequency spectrum over time

The expected histogram of mutation frequencies in a sample (the “site frequency spectrum”) is a geometrically decreasing function of increasing mutation frequency under the standard neutral model (Wakeley, 2008). Departures from this expectation are often summarized as single numbers whose expectations are ≈ 0 under this null model. In this section, I describe the average dynamics of two widely used statistics (Tajima, 1989; Zeng *et al.*, 2006) as a function of both time since the optimum shift and of distance from trait-affecting mutations.

Figure 8 shows the average behavior of Tajima’s (1989) *D* over time. Figure 8a shows the mean *D* per window, averaging across loci and across replicates. Prior to the optimum shift, the mean *D* is negative in the central window containing selected variants. For highly polygenic traits, the equilibrium *D* is *≈ -*0.1 in this window due to a large number of rare deleterious alleles segregating. After the optimum shift, *D* becomes more negative when the optimum shift is large and the mutation rate is smaller. In linked windows, the magnitude of the change in averaged *D* decays rapidly with increasing genetic distance.

**Figure 8:**
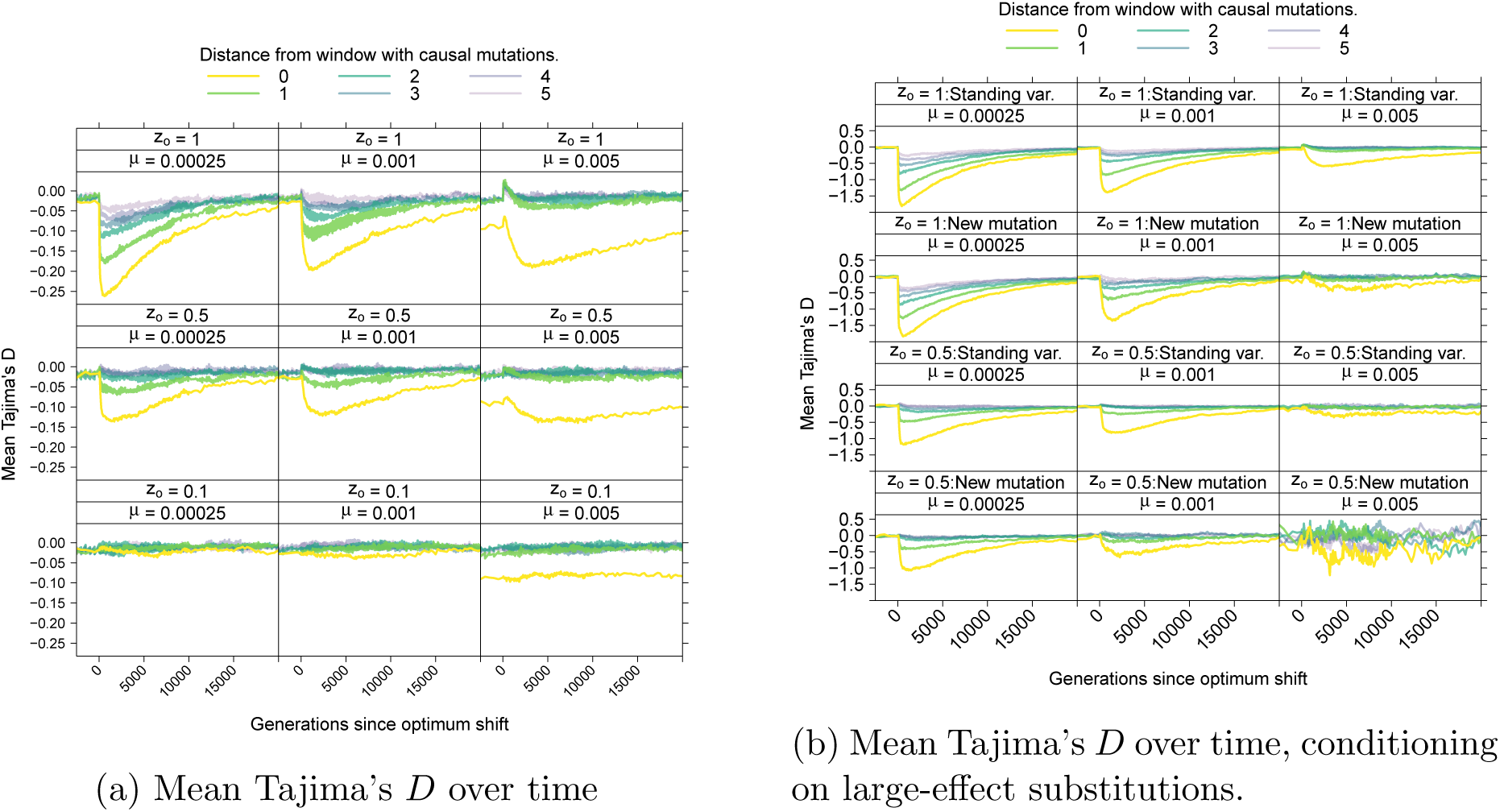
Tajima’s (1989) *D* statistic over time. 8a shows the average value of Tajima’s (1989) *D* over time. The data are shown separately for windows of different distances from the central window where mutations affecting the trait arise (Figure 2) 8b shows the mean value of *D* conditioning on a locus fixing a mutation with effect size *Nγ*^2^ ≥ 100. These loci are separated by whether the fixation was from standing variation, meaning a mutation predating the optimum shift, or from a new mutation arising after the shift.

Averaging over loci experiencing large-effect fixations, Figure 8b shows a stronger hitch-hiking pattern centered on the window containing selected variants. Although the deviation in *D* from equilibrium decays relatively quickly both along a chromosome and over time, large-effect substitutions generate sufficiently negative *D* values that such loci will be enriched in the tails of empirical distributions of the statistic. Qualitatively similar patterns hold for the overall reduction in diversity (Figure S7) and the *H′* statistic (Figure S8). The latter statistic returns to equilibrium rather rapidly, consistent with previous results (Przeworski, 2002).

Here, large-effect fixations from new mutations and from standing variants have similar average effects on statistics like *D* and *H′* (Figures 8 and S8). Figure 9 shows the number of haplotypes at a locus associated with sweeps from standing variation as a function of the effect size of the variant. Here, a haplotype is defined as a unique genotype at a locus, including all neutral and non-neutral variants. Large-effect sweeps from standing variation are either extremely rare (at high *µ*) or are rare at the time of the optimum shift when *µ* is small, and usually associated with very few, and often only one, haplotype at the onset of directional selection (Figure 9).

**Figure 9:**
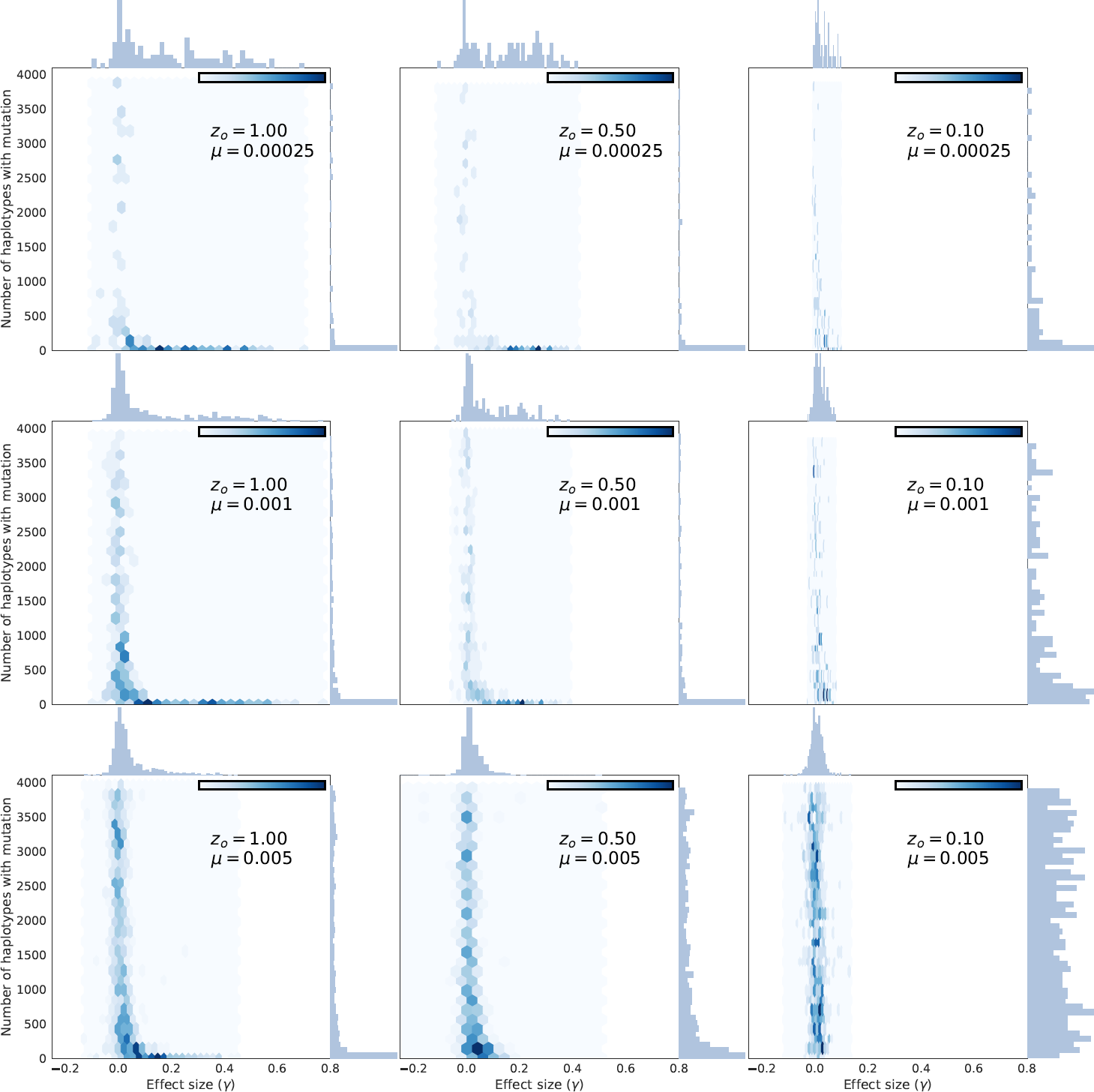
The number of haplotypes associated with fixations from standing variation of different effect sizes. Each panel shows the effect size of a fixation from standing variation (x axis) and the number of unique haplotypes in the entire population containing that mutation. The number of haplotypes for each mutation is taken immediately prior to the optimum shift and excludes any mutations that arose that generation. Thus, all mutations found on a single haplotype are more than one generation old.

#### Power to reject the null model using the site-frequency spectrum

Figure S9a shows the power to detect a value of *D* more negative than expected under the standard neutral model, after applying a multiple testing correction such that the per-window rejection rate under the null model is 0.05. The overall power of the test is low due to the number of tests performed (one per window), and is consistent with previous work (Braverman *et al.*, 1995; Przeworski, 2002). However, the set of loci representing “significant” deviations from the null model are enriched for large-effect substitutions (Figure S9b), of which there are relatively few per replicate (Figure S10). When mutation rates are smaller, “significant” *D* values are most common at loci where large-effect mutations fixed. As the trait becomes more polygenic and/or the optimum shift less drastic, the enrichment shifts towards sweeps from standing variation.

The behavior of *H′* is similar to that of *D*, but power decreases more rapidly with time since the optimum shift (Figure S11a, (also see Przeworski, 2002)). The behavior of a related test, the compsite likelihood ratio test of (Nielsen *et al.*, 2005), evaluated using SweeD (Pavlidis *et al.*, 2013), is qualitatively similar to that of *H′* (Figure Figure S12).

#### Haplotype homozygosity

Rapid increases in allele frequency due to selection will result in long stretches of homozygosity flanking the selected mutation (Kim and Nielsen, 2004). Summaries of haplotype homozygosity are widely used to detect recent selection (Voight *et al.*, 2006; Ferrer-Admetlla *et al.*, 2014) and are indirect summaries of the underlying linkage disequilibrium in the data (Sabatti and Risch, 2002).

The *nS*_*L*_ statistic (Ferrer-Admetlla *et al.*, 2014) measures the ratio of homozygosity on the ancestral allele to that on the derived allele for each variant in the data. A negative value of the statistic implies longer runs of homozygosity around the derived allele. Figure S13 shows the average behavior of z-scores obtained from binning *nS*_*L*_ scores by derived allele frequency (see Methods). The signal of strong positive selection, indicated by a negative z-score, is short-lived and only observed when the mutation rate is smaller and the optimum shift is large. The signal is also restricted to regions very near where selected mutations arise.

Shortly after the optimum shift, the mean *z*-score becomes positive (Figure S13). This temporal dynamic is qualitatively similar to what is seen under standard models of selective sweeps, as the time since the sweep moves further into the past (Figure S14) Thus, the positive z-scores in Figure S13 may be interpreted as either older sweeps from new mutations or strong sweeps from common variants. However, the latter class of sweeps does not occur in these simulations (Figure 9). This difficulty in interpretation is a general issue arising from the fact that patterns of variation due to strong sweeps from standing variation overlap considerably with those of older sweeps from new mutations (Schrider *et al.*, 2015).

A related class of statistics designed to detect strong sweeps from standing variation are based on the overall haplotype diversity in a window (Garud *et al.*, 2015). The temporal patterns associated with these statistics are again short-lived and are all in the direction of reduced overall haplotype heterozygosity, which is a signal of strong sweeps from new mutations (Figures S15, S16, and S17).

### Robustness to variation in the recombination rate

In this section, I explore the effect of varying the scaled recombination rate within a locus, *ρ*. At higher mutation rates, longer fixation times are more likely as *ρ* decreases (Figure 10). In individual replicates, there is a tendency towards negative disequilibrium among beneficial mutations (*γ >* 0, Figure S18), suggesting a role for interference among selected sites affecting times to fixation (Hill and Robertson, 1966; Felsenstein, 1974). In the previous sections, the ratio of *ρ* to *θ* within loci is one, which is roughly “human”-like (Dumont and Payseur, 2008; SÉgurel *et al.*, 2014). For species like *Drosophila melanogaster*, where *ρ* ≫ *θ* (Haddrill *et al.*, 2005), fixation times will be much shorter on average (Figure 10). Note that the effect of recombination rate on fixation time is most dramatic for *µ* = 0.005, which is also the part of the parameter space explored here where fixations of larger-effect (*Nγ*^2^ ≥ 1, 000) are rare.

**Figure 10:**
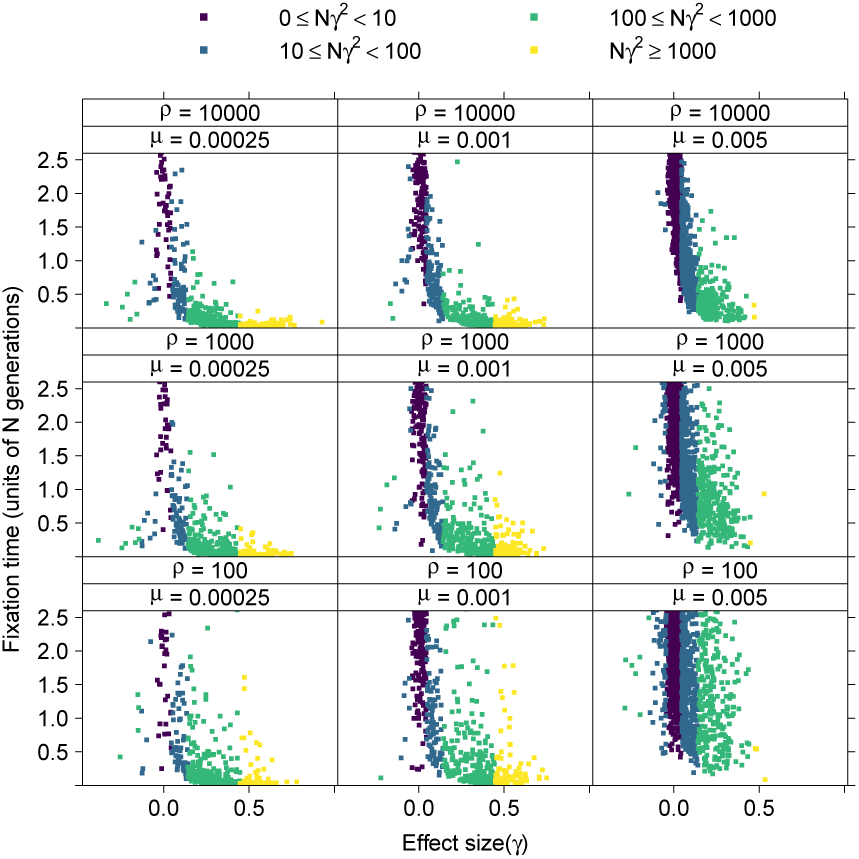
The sojourn times of fixations in a ten locus system with varying recombination rates within loci. The x axis is the effect size of a fixation, and the y axis is its time to fixation scaled by the population size. The expected fixation time due to drift alone is 4 and the distribution of fixation times under neutrality has a long tail including large values. The points are collected from 256 replicates for each parameter combination and colored by order-of-magnitude ranges of their scaled strength of selection (*Nγ*^2^).

The within-locus recombination rate has no discernible average effect on 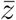 nor on *V*_*G*_ (Figure S19). The differences in the height of the “spike” in *V*_*G*_ when *µ* = 2.5 × 10^−4^ show no clear pattern with *ρ* and are thus attributable to Monte Carlo error in estimating a second-order statistic from 256 replicates.

Unlike the mean trait value and variance, the mean temporal dynamics of summaries of variation data are strongly affected by *ρ* (Figure S20) as expected Kaplan *et al.* (1989); Braverman *et al.* (1995). Figure S21 shows how the within-locus recombination rate affects patterns of haplotype diversity in a ten locus system with *σ*_*γ*_ = 0.25. When *ρ* is small, the impact of linked selection is much more apparent. These effects of the local recombination rate on patterns of hitch-hiking are expected from standard theory of directional selection because both the magnitude and extent along the genome of linked selection depends on the ratio of the recombination rate to the selection coefficient (Kaplan *et al.*, 1989; Durrett and Schweinsberg, 2004; Nielsen *et al.*, 2005).

### Varying the distribution of effect sizes

The results described in the previous sections are based on a Gaussian distribution of effect sizes whose standard deviation is held constant. In this section, I vary the distribution of effect sizes (DES) such that the fraction of mutations with *Nγ*^2^ ≥ 100 varies and compare the average dynamics of adaptation and patterns of hitch-hiking. I also compare a Gaussian distribution to a Gamma distribution with different shape parameters. In order to simplify the presentation, I only show results for the case of a large optimum shift (*z*_*o*_ = 1), which is the case resulting in the most extreme hitch-hiking signals. I compare the results of Gaussian distributions of effect sizes to two gamma distributions with shape parameters of one and one-half.

Varying the fraction of large-effect mutations has a weak effect on the mean time to reach the new optimum, with traits with low mutation rates adapting more slowly on average when the majority of variants are of small effect (Figure S22a). This observation should be unsurprising as the population must wait longer for a strongly selected mutation in this case.

Patterns of variation expected due to hitch-hiking are more extreme when *Pr*(*Nγ*^2^ ≥ 100) is small, as the population has to wait longer for strongly selected variants (Figure S23). The overall pattern is that the average differences between DES are subtle, with Gamma distributions showing less extreme hitch-hiking patterns (negative values) on average than the Gaussian DES. However, this difference between DES is only observed when both the mutation rate and the proportion of new mutations of large effect are both small.

## Discussion

I have used simulations to describe the average behavior of selected and neutral mutations during the adaptation of a quantitative trait to a single, sudden shift in the optimal trait value. The genotype-to-phenotype model considered here is the classic model of evolutionary quantitative genetics, assuming strictly additive mutational effects on trait value with fitness determined by Gaussian stabilizing selection (Turelli, 1984; Barton, 1986; BÜrger, 2000). The primary goal here was to merge this model of a phenotype with the simulation methods commonly used in population genetics to study the effect of natural selection on the dynamics of linked neutral variation (Kaplan *et al.*, 1988, 1989; Braverman *et al.*, 1995; Przeworski, 2002; Innan and Kim, 2004).

The simulations performed here have several important differences from recent theoretical treatments of adaptation to sudden optimum shifts (see below). However, the conditions for a selective sweep are consistent with predictions made using theoretical results from from Jain and Stephan (2017b) and HÖllinger *et al.* (2019). A direct comparison to the quantitative predictions from Jain and Stephan (2017b) is difficult because their expressions depend on the assumption of equal forwards and backwards mutation rates at each position. However, several qualitative comparisons can be made. First, the simulations presented here are comparable to the “most effects are large” case from Jain and Stephan (2017b) because the trait variance *increases* during adaptation (also see de Vladar and Barton (2014)) due to large-effect mutations moving from low to intermediate frequency. Mutations with large effects on trait values at the time of the optimum shift are most likely to rise in frequency (Figure 4 and 5), although mutations that eventually fix are not necessarily those with the largest effect size. When several large-effect mutations co-segregate, those with the highest initial frequencies tend to reach fixation. If initial frequencies are similar, the variant with the highest initial fitness typically fixes. For a given distribution of effect sizes, faster sweeps are more likely at lower mutation rates.

Regimes where the genetic variance decreases during adaptation are not possible for any of the simulations presented here. The decrease in variance is seen in the “most effects are small” domain where the equilibrium frequency of variants prior to the optimum shift is one-half, which maximizes the variance (de Vladar and Barton, 2014; Jain and Stephan, 2017b). Adaptation to the new optimum displaces allele frequencies, reducing the variance from its maximum value (see, for example Figure 9 of de Vladar and Barton (2014)). However, the equilibrium frequency of one-half for small effect mutations requires equal rates of forward and back mutation de Vladar and Barton (2014); Jain and Stephan (2017b), and is therefore incompatible with the infinitely many sites assumption made here.

When considering the pattern of hitch-hiking at a locus, the presence or absence of a large-effect fixation at a locus is a reliable predictor of the magnitude of hitch-hiking patterns. As expected, such fixations are more common when the mutation rate is smaller (HÖllinger *et al.*, 2019) and thus strong departures from equilibrium patterns of variation are not expected for more polygenic traits (Figure 8). For the optimum shift model considered here, the strength of selection is not constant over time (Figure 5, see also Kimura (1981)). Thus, genotypes containing variants that were initially strongly favored by selection are much more weakly selected by the time the population has reached the new optimum. This weakening of selection increases fixation times to the order of the population size (Figure S2), which is much longer than the times *" N* generations expected for directional selection in large populations (Stephan *et al.*, 1992).

The exploration of hitch-hiking signals here involved simulation ten unlinked loci within which mutations affecting the trait were concentrated in a central window (Figure 2). While the ratio of recombination to mutation events is at least nine to one for the majority of the results shown here (see Methods), it is possible that signals of selection are made more pronounced by to the localization of selected mutations and should be explored further.

Here, the number of selected mutations segregating over time ranged from dozens to several hundred, as a function of the underlying mutation rate (Figure S24). At high mutation rates, the number of segregating loci are roughly the same as some of the results presented in de Vladar and Barton (2014). Here, however, the partial linkage among sites in this work leads to some negative linkage disequilibrium (Figure S18), which is a signal of interference (Hill and Robertson, 1966; Felsenstein, 1974). This interference has little effect on the mean time to adaptation, but fixation times are increased. The lack of effect on time to adaptation is driven by initial large fitness differences among genotypes (Figure 5, also see HÖllinger *et al.* (2019)). Once the population is close to the new optimum, selection on individual genotypes is much weaker (Figure 5), setting up the conditions for interference to affect fixation times (Hill and Robertson, 1966).

The distribution of effect sizes has different effects on properties of the trait than on patterns of hitch-hiking. The mutation rates used here span the parameter space from partial and complete sweeps being most common to the optimum being reached via allele frequency shifts of many mutations (Figure 4, HÖllinger *et al.* (2019)). In general, the mean time to adapt is not strongly affected by the DES if the fraction of new mutations of large effect is constant (Figure S22a). For a given mutation rate, lowering the mutational variance lowers the probability of a strongly selected mutation, resulting in stronger signals of hitch-hiking (Figure S23). Effectively, the trait becomes more mutation limited when the mutational variance is lower, although the effect is most pronounced when the mutation rate itself is small (Figure S23). When the trait is more polygenic, the average patterns of variation are not strongly dependent on the DES nor on the proportion of new variants with large effect (Figure S23).

The genetic model assumed here does not lead to sweeps of large-effect mutations from common variants (frequencies greater than, say, five percent). Rather, the stabilizing selection around the initial optimum keeps large-effect mutations rare, such that sweeps from such standing variations start at low frequencies. Importantly, it is not possible to tune the model parameters to obtain sweeps from large-effect, but common, variants with high probability. Changing the strength of stabilizing selection (*V*_*S*_) preserves the rank orders of fitness for all genotypes, merely changing how fit they are in an absolute sense. One could randomly reassign effect sizes at the time of the optimum shift in an attempt to approximate a gene by environment interaction. However, such a procedure would be arbitrary, and thus not represent a principled model for generating *detectable* “soft sweep” patterns. Rather, it is tempting to invoke a need for pleiotropic effects in order to have large-effect mutations segregating at intermediate frequencies at the time of the optimum shift, with the shift itself accompanied by a change in the covariance between trait values and fitness.

It is important to note a key methodological difference between this work and that of other authors. HÖllinger *et al.* (2019) stopped their simulations when the population was close to the new optimum while the simulations conducted here allowed evolution to continue much longer. Thus, on the time scale during which the population adapts, fixations are not observed when Θ is high (see Figure 4 of HÖllinger *et al.* (2019)). Here, we observe fixations of large effect for the mutation rates corresponding to Θ = 4*Nµ* = 100 (Figure 4), which HÖllinger *et al.* (2019) show is the parameter range where adaptation occurs primarily by changes in allele frequency. These results are consistent with the theoretical predictions from HÖllinger *et al.* (2019), as the fixations in the simulations described here take place on time scales longer than the mean time to reach the new optimum. In the rightmost column of Figure 4, the population has adapted quickly, with the fixations occurring over a much longer time scale (Figure S1). Likewise, the leftmost column of Figure 4 corresponds to Θ = 5, where we observe a mixture of partial and complete selective sweeps by the time the new optimum is reached, which is expected from the theory presented in HÖllinger *et al.* (2019).

This work (and that of HÖllinger *et al.* (2019)) differs from the analytical and numerical work of de Vladar and Barton (2014); Jain and Stephan (2015, 2017b) in several key aspects. First, we consider irreversible mutation here (the infinitely many sites model of Kimura (1969)), while de Vladar and Barton (2014) assumed equal rates of forward and reverse mutation (see also Barton (1986); Jain and Stephan (2015, 2017b)). The infinitely many sites model used here was chosen because it is the most commonly used mutational model for investigating the effects of linked selection during adaptation (*e.g.* Braverman *et al.*, 1995; Przeworski, 2002; Przeworski *et al.*, 2005). I also allow for partial linkage among sites, which is a key difference from the work based on the Barton (1986) framework, which assumes free recombination. As noted above, partial linkage affects the long-term dynamics of selected mutations (Figure 10).

I have focused on standard summaries of variation data that have widely been applied to detect selection from sequence data. The behavior of the majority of such summary statistics have only been tested using coalescent simulations of strong selection on a single sweeping variant, which is the dominant generative model used for making predictions about linked selection. Thus, it is unsurprising that these statistics show the strongest departures from equilibrium neutrality for traits with low mutation rates. However, an important observation here is that the mean behavior of these statistics are similar for sweeps from new mutations and sweeps from standing genetic variation, which is a consequence of the standing variants being rare at the onset of selection (Figure S6, also see Berg and Coop (2015); Przeworski *et al.* (2005); Orr and Betancourt (2001); Hermisson and Pennings (2005)). The only test statistic based on patterns of SNP variation for detecting polygenic adaptation that I am aware of is the singleton density score (Field *et al.*, 2016). I have not explored this statistic here, as it would be more fruitful to do so using simulations of much larger genomic regions applying tree sequence recording (Kelleher *et al.*, 2018) and explicit modeling of trait architectures at nor near the infinitesimal limit Robertson (1970, 1977). It also appears that the magnitude of selective effects on phenotypes attributable to changes in the singleton density by Field *et al.* (2016) were substantially overestimated due to uncontrolled-for population structure in the GWAS data, and there is little evidence for selection on height when the analysis was redone using effect sizes from the UK Biobank data (Berg *et al.*, 2019; Sohail *et al.*, 2019).

I have only considered the equilibrium Wright-Fisher model here. However, it is well-understood that departures from this demographic model affect patterns of neutral variation and thus the detection of regions affected by linked selection (Thornton and Andolfatto, 2006; Thornton and Jensen, 2007; Thornton *et al.*, 2007; Jensen *et al.*, 2007, 2008). Demographic departures from constant population size indeed affect the prevalence of sweeps and the rate of phenotypic adaptation in optimum shift models (Stetter *et al.*, 2018). Here, we are primarily interested in how the parameters affecting the trait’s “architecture”, mainly the parameters affecting the mutational variance of the trait, impact patterns of linked selection.

It is crucial to restate the assumptions of the genetic model assumed here, which involves strictly additive effects on a single trait under “real” stabilizing selection (Johnson and Barton, 2005). This model is the standard model of evolutionary quantitative genetics (Turelli, 1984; Barton, 1986; BÜrger, 2000), which is why it is the focus of this work. However, a more thorough understanding of the dynamics of linked selection during polygenic adaptation will require investigation of models with pleiotropic effects (*e.g.*, Zhang and Hill, 2002; Simons *et al.*, 2018). Because the adaptation to the new optimum is rapid when the mutation rate is large, the allele frequency changes involved are also small when mutational effects are pleiotropic (Simons *et al.*, 2018). The question in a pleiotropic model is the role that large-effect mutations may play, which is an unresolved question.

The simulations here also model the entirety of heritable variation for the trait. An alternative approach would be to allow for a unlinked additive genetic background with its own mutational variance. Such an approach would be straightforward assuming an infinitesimal model for the background, as has been done previously (Chevin and Hospital, 2008; Stetter *et al.*, 2018). Stetter *et al.* (2018) simulated “domestication” traits evolving to a new optimum via truncation selection and a heritable background affecting the focal trait. They concluded that the contribution of genetic background to several outcomes of interest (speed of adaptation, fixations of beneficial mutations, etc.) was of overall less importance to the dynamics than the variance in mutational effect sizes, *σ*_*γ*_. Clearly, however, the details will depend on the specifics of the model, with Chevin and Hospital (2008) at one extreme and the current work at perhaps the other. Here, the simulations with high mutation rates imply that any single segregating variant finds itself in a mutation-rich genetic background of up to several hundred segregating variants, the majority of which have small fitness effects (Figure S24). Another appealing alternative would be to simulate entire genomes using an adaptation of Robertson’s (1977) method to incorporate tree sequence recording (Kelleher *et al.*, 2018) and large-effect mutations occurring at some rate. Such a scheme would generate large-effect genomic regions through two different mechanisms: the occasional large-effect mutation as well as via large-effect haplotypes arising from stochastic recombination events (Sachdeva and Barton, 2018).

It may also be of interest to explore non-additive genetic models in future work. In particular, models of non-complementing recessive effects within genes are a specific class of models with epistasis that deserve consideration due to their connection with observations of allelic heterogeneity underlying variation in complex traits (Clark, 1998; Thornton *et al.*, 2013; Sanjak *et al.*, 2017; Gruber and Long, 2009; King *et al.*, 2014; Long *et al.*, 2014; McClellan and King, 2010; Chakraborty *et al.*, 2018). Acknowledging the focus on the standard additive model, the current work is best viewed as an investigation of a central concern in molecular population genetics (the effect of natural selection on linked neutral variation) having replaced the standard model of that sub-discipline with the standard model of evolutionary quantitative genetics. As laid out by several authors (Jain and Stephan, 2017b,a; Messer and Petrov, 2013), there are considerable theoretical and empirical challenges remaining in the understanding of the genetics of rapid adaptation. For models of phenotypic adaptation, our standard “tests of selection” are likely to fail, and are highly under-powered even when the assumptions of the phenotype model are closer to that of the standard model.

## Acknowledgments

The author thanks Jaleal Sanjak, David Lawrie, Jeffrey Ross-Ibarra, Markus Stetter, Tony Long, Joachim Hermisson, Nick Barton, Kavita Jain, and two anonymous reviewers for valuable discussions and feedback on the manuscript. This work is supported by NIH Grant R01-GM115564 to the author.

## Supporting Information

**Figure S1:**
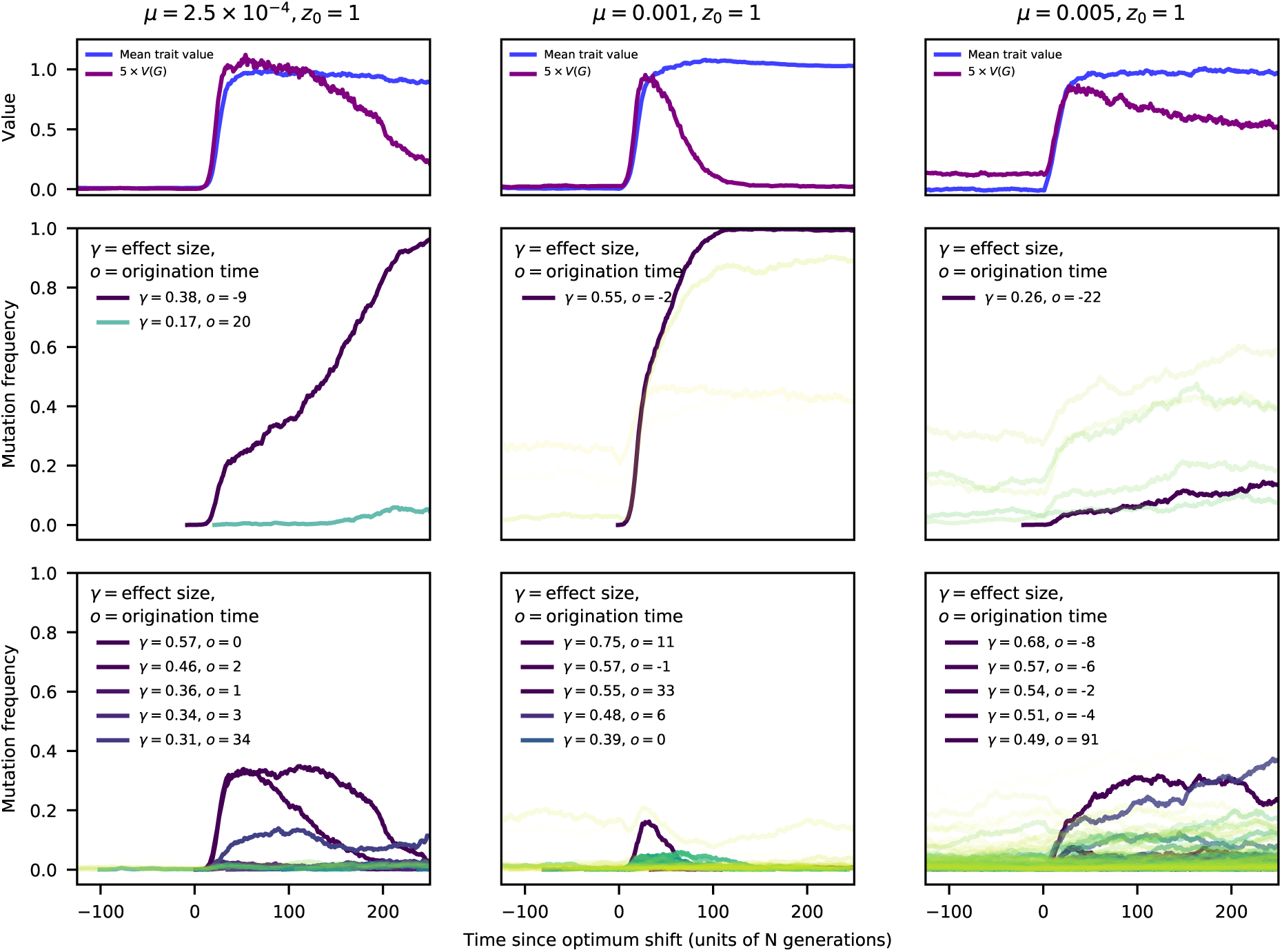
Trajectories of selected mutations, shown over a short time scale. The data and colors are identical to Figure 4.

**Figure S2:**
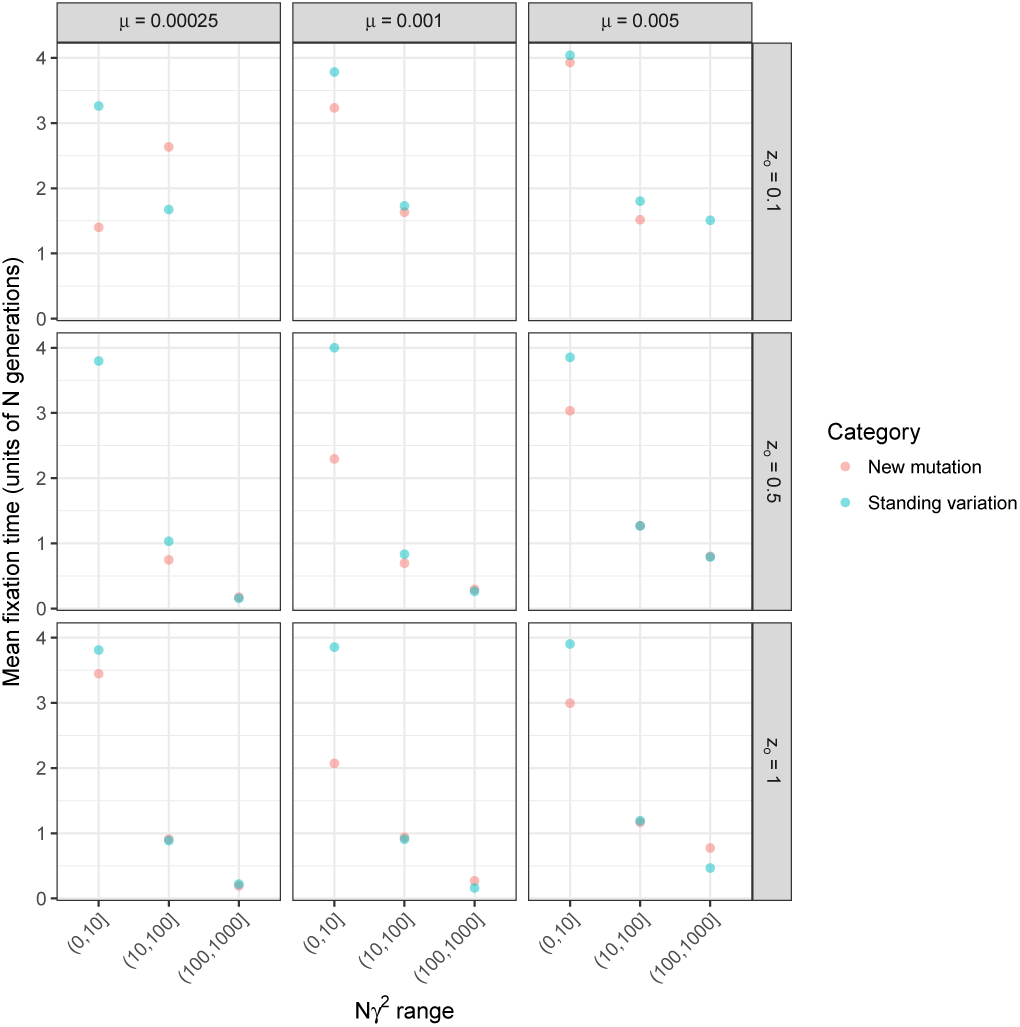
The mean time to fixation of a selected mutation in a single-region simulation. The mean times are scaled by population size (*N*) to facilitate comparison to the expectation in the absence of selection, which is 4. The mean values are separated by both scaled strength of selection (*Nγ*^2^) and by fixations from standing variation versus new mutations arising in the first one hundred generations after the optimum shift.

**Figure S3:**
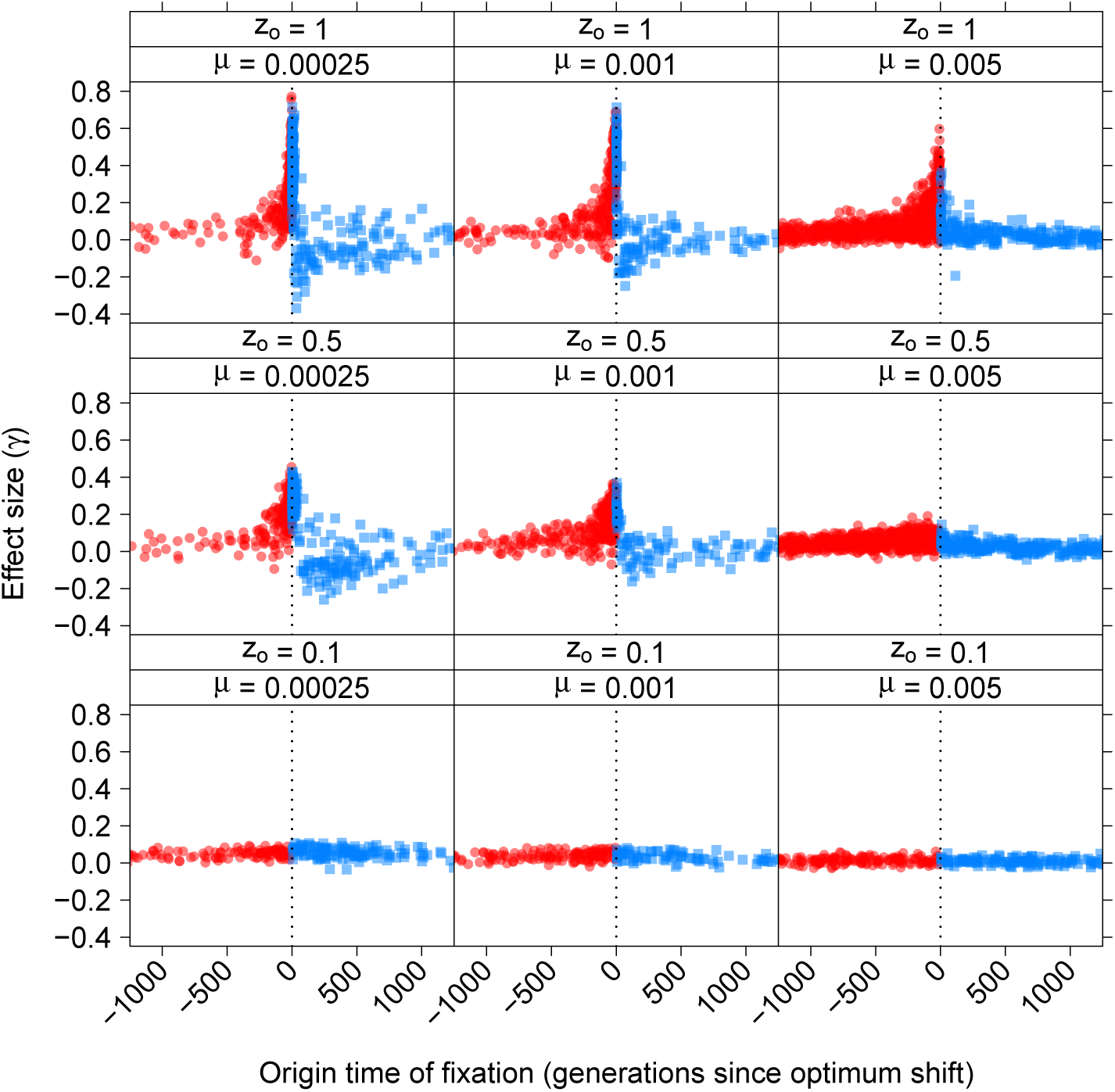
Fixations with large effects on trait values typically arise close to the time of the optimum shift. The x axis shows the origin time of each fixation and the y axis is that fixation’s effect size, *γ*. Time is measured in units of *N* generations an scaled so that the optimum shift occurred at time zero. Red points are origin times prior to the optimum shift and are therefore fixations arising from standing variation. Blue points are fixations from new mutations.

**Figure S4:**
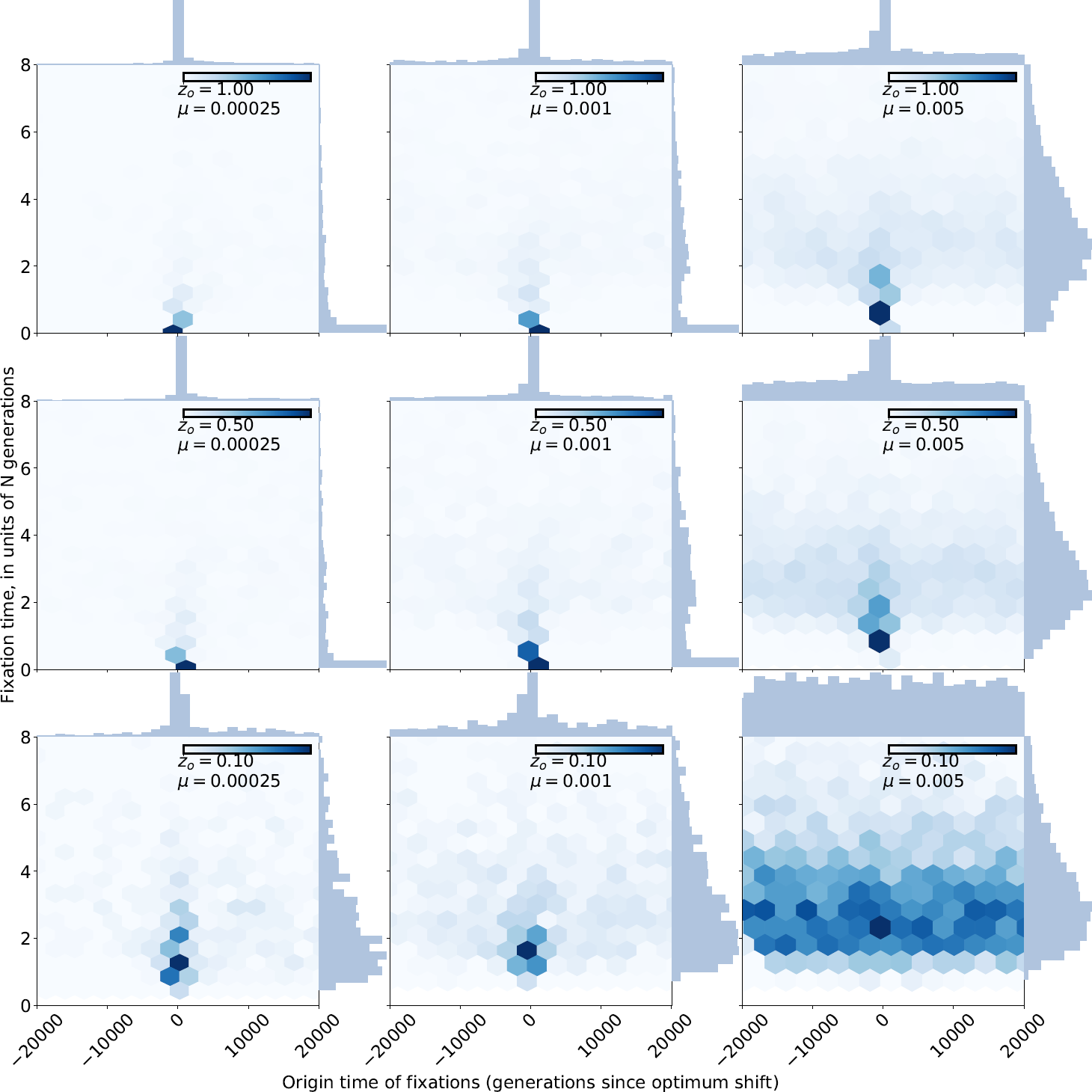
The time to fixation as a function of origin time. The x axis shows the origin time of each fixation in units of *N* generations and scaled such that the optimum shift is at time zero. The y axis is the fixation time, again in units of *N* generations. Each panel shows the joint density of origin and sojourn time as well as the marginal distributions, and the raw data are the same as in Figure S3. As the trait is more mutation-limited (smaller *µ*) and/or the optimum shift more extreme (larger *z*_*o*_), the fixation time decreases because the effect sizes increase (Figure S3).

**Figure S5:**
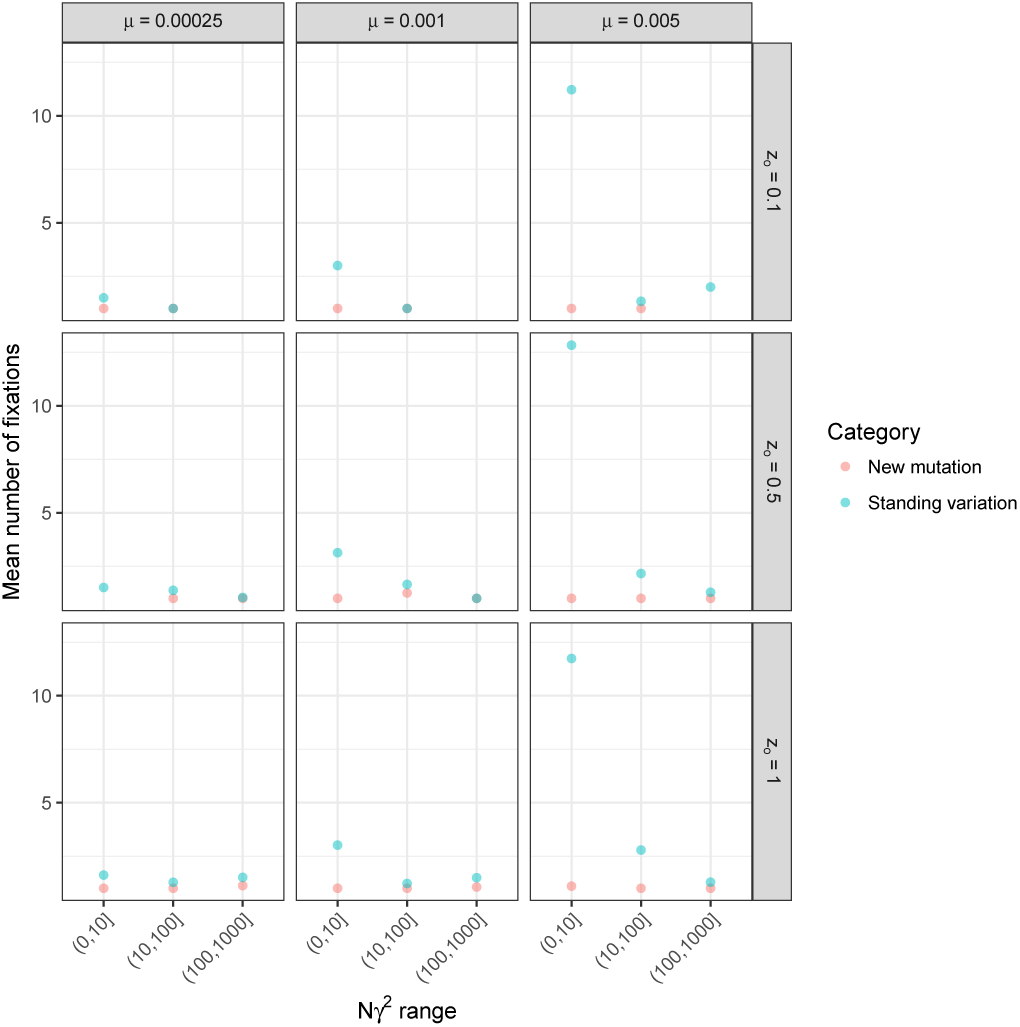
The mean number to fixations of selected mutations in a single-region simulation. The mean values are separated by both scaled strength of selection (*Nγ*^2^) and by fixations from standing variation versus new mutations arising in the first one hundred generations after the optimum shift.

**Figure S6:**
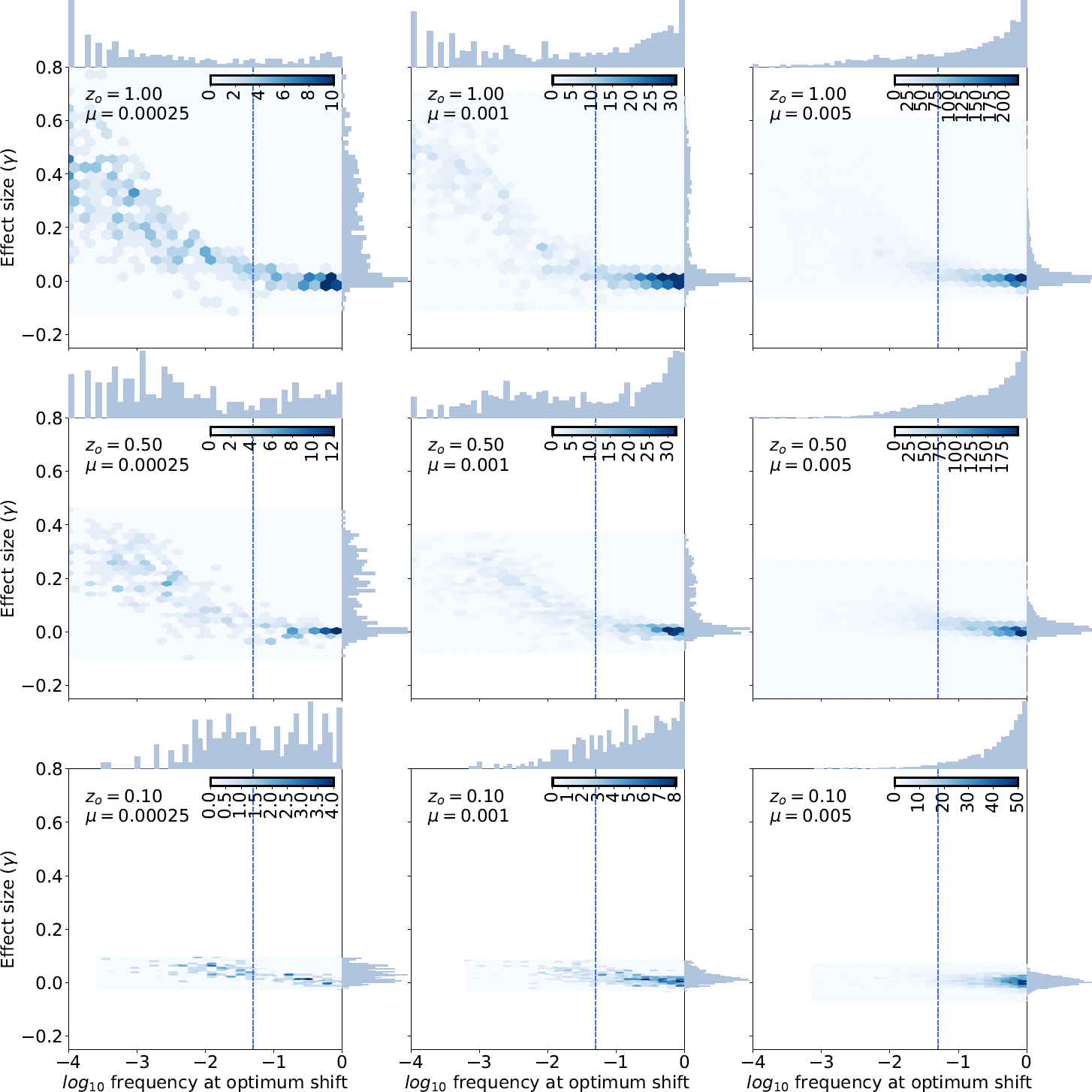
The joint distribution of frequency at the time of the optimum shift and effect size (*γ*) for fixations from standing variation. Each panel shows the joint density of the mutation frequency at the time of the optimum shift and effect size, using the raw data from Figure S3 The vertical dashed line represents a frequency of 0.05, which Przeworski *et al.* (2005) identified by simulation as a rough threshold below which sweeps from standing variation will show a reduction in levels of neutral sequence diversity at linked sites, provided that they are associated with large effect on fitness. As the trait is more mutation-limited (smaller *µ*) and/or the optimum shift more extreme (*z*_*o*_ larger), there is more of a tendency for sweeps from standing variation to be chosen from rarer mutations with larger effects on trait values. Sweeps from common variants are associated with small effect sizes.

**Figure S7:**
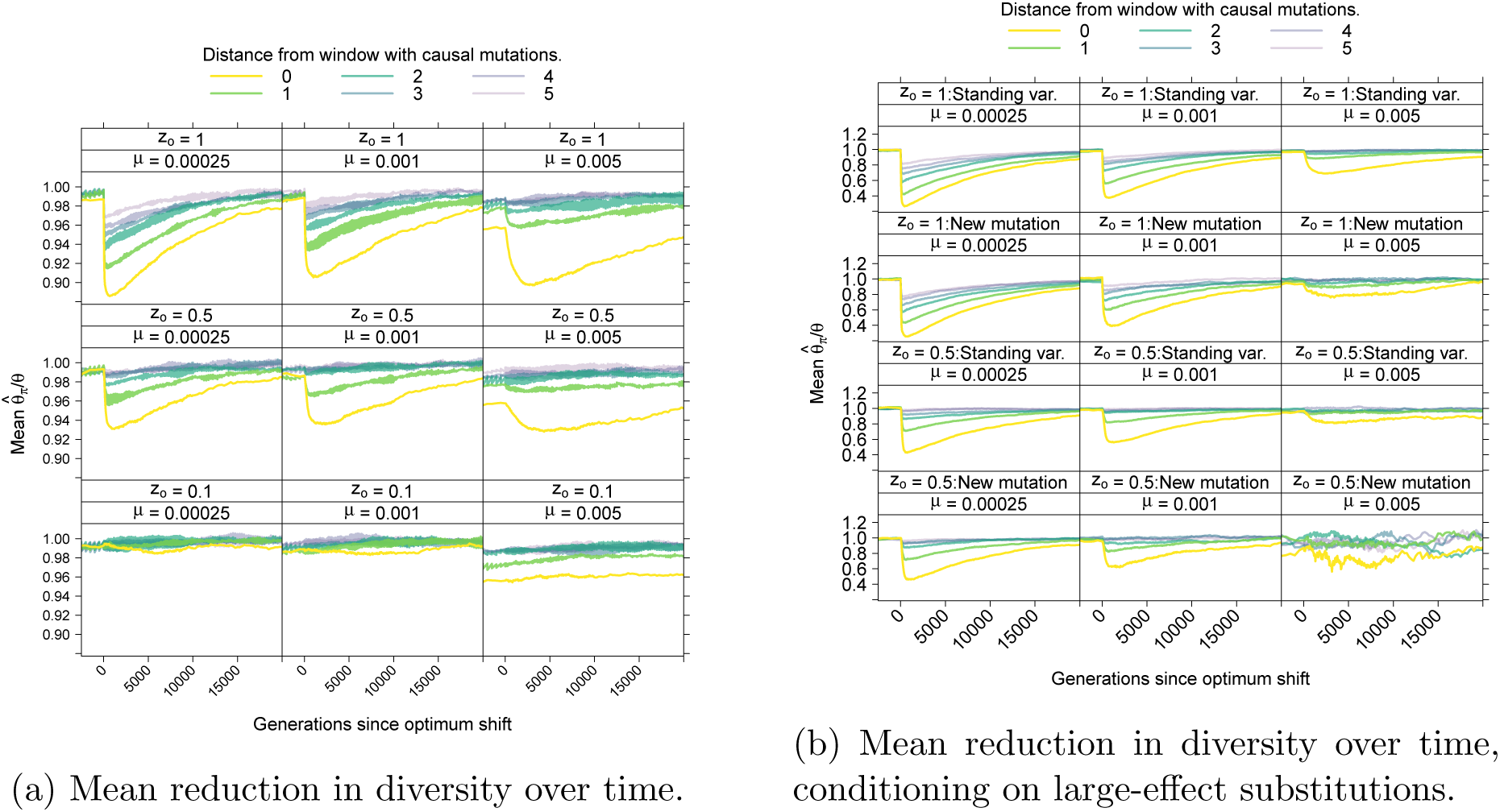
The average reduction in genetic diversity due to the optimum shift. Here, diversity is measured as the mean number of pairwise differences between sequences (Tajima, 1983) divided by *θ* = 4*N*_*e*_*µ*. S7a shows overall reduction in diversity, averaging across all ten loci and across all replicates. The mean value is shown separately for windows of increasing distance from the central window containing selected mutations (Figure 2). A distance of zero refers to the central window. There are an equal number of flanking windows on either side of the central window (Figure 2) and therefore a distance of one averages the results for the windows immediately to the left and right of the central window and a distance of two is averaging the next window on either side, etc.. S7b shows the mean reduction in diversity at loci experiencing a sweep due to a mutation with effect size *Nγ*^2^ ≥ 100. Sweeps from standing variants (those arising prior to the optimum shift) and new mutations arising after the shift are shown separately. There are no lines for the lower right panel because no large-effect fixations occur in that part of the parameter space. No panels are shown for an optimum trait value of *z*_*o*_ = 0.1 for the same reason.

**Figure S8:**
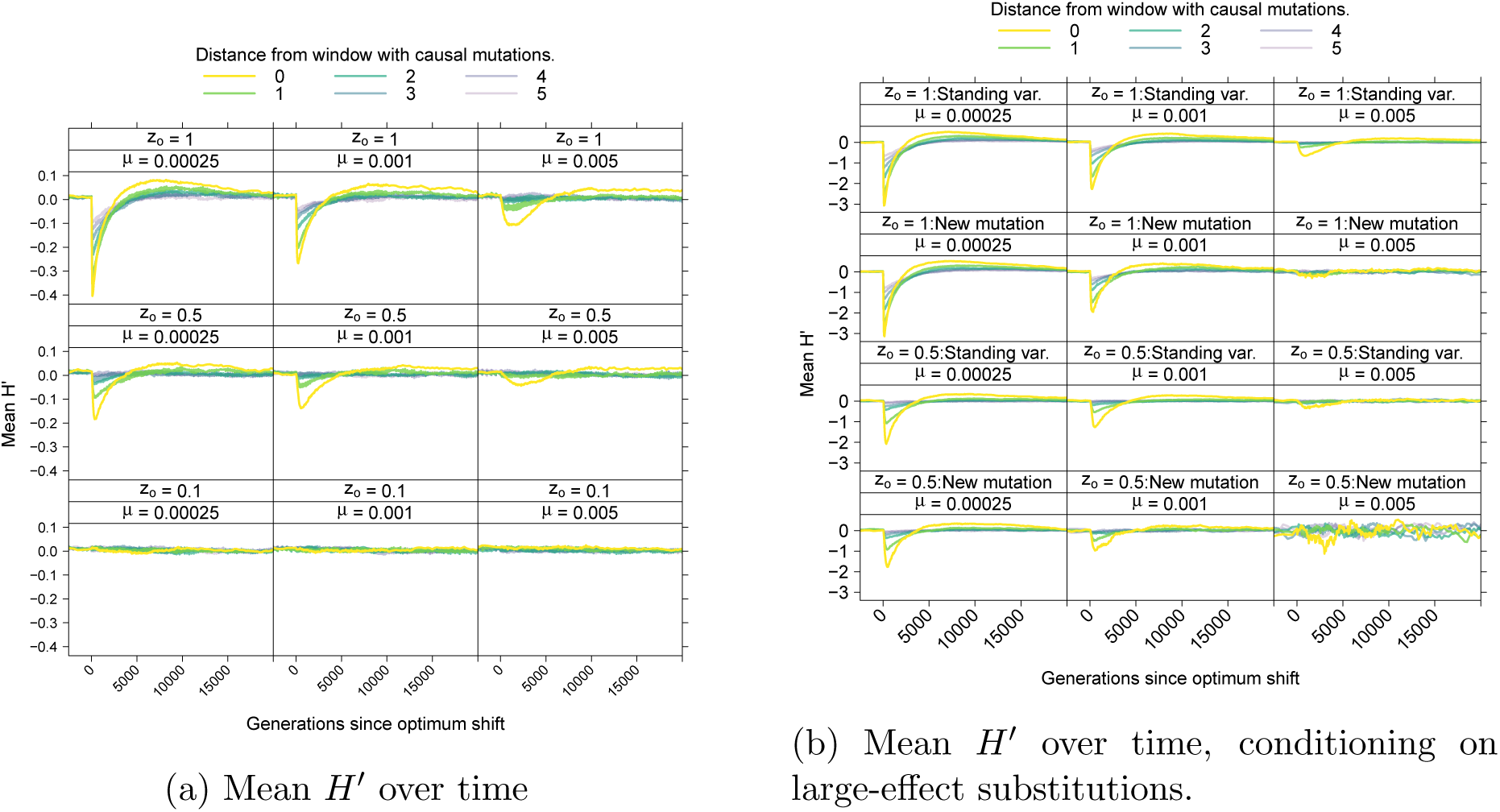
Average behavior of *H′* (Zeng *et al.*, 2006) over time. S8a shows the mean *H′* over time, averaging over both loci and over replicates. The data are plotted as a function of increasing window distance from where mutations affecting the trait occur (Figure 2). S8b shows the mean of *H′* over time, conditioning on the fixation of a large-effect mutation either from standing variation or from a new mutation arising after the optimum shift.

**Figure S9:**
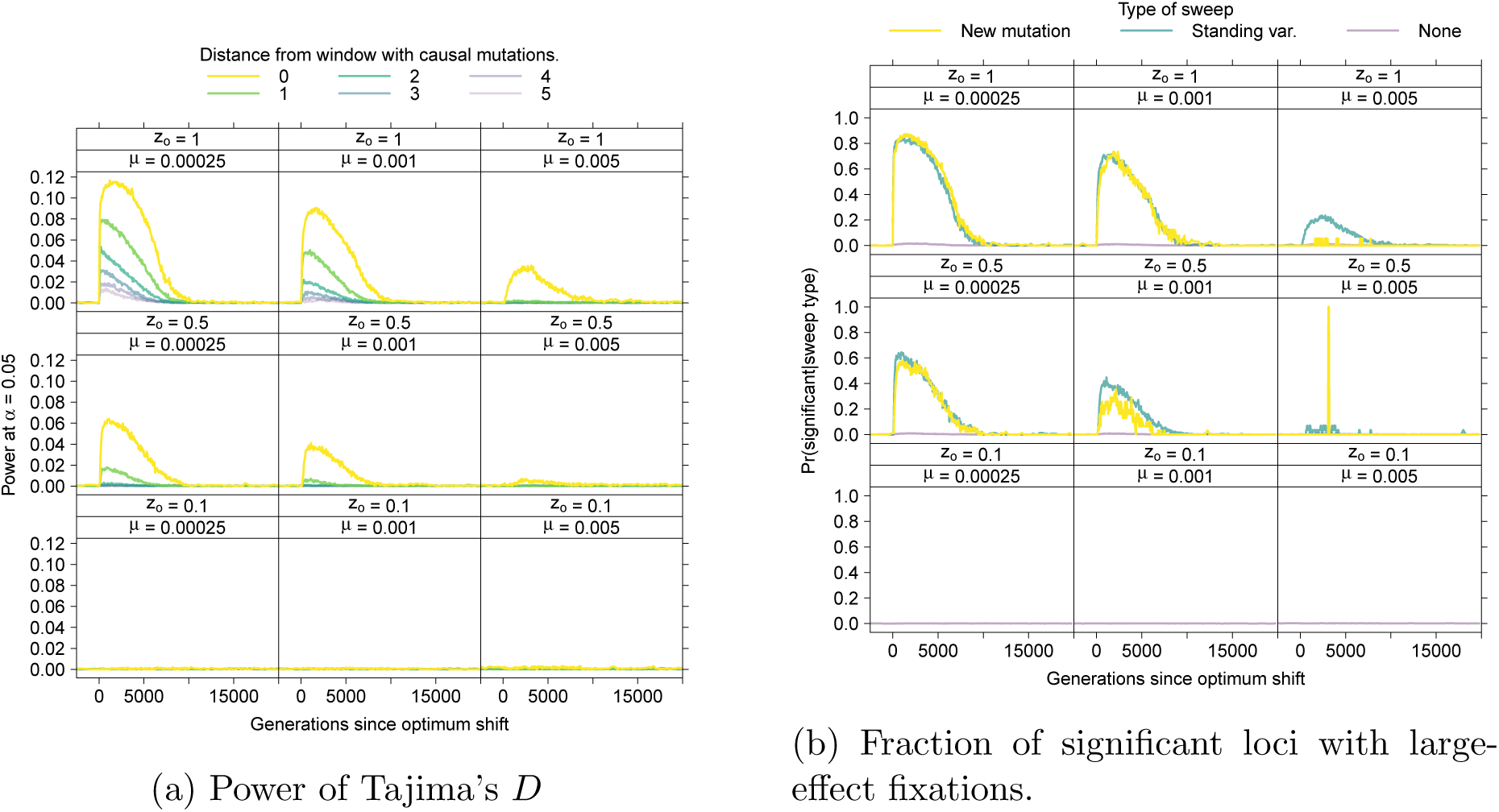
Power of Tajima’s (1989) to reject the standard neutral model. S9a The power to reject the null model in windows of increasing distance from selected site as a function of time since the optimum shift, corrected so that the per-window rejection rate under the null model is 0.05. The windows are labeled using the same scheme as Figure 8. S9b The probability of a “significant” window at a locus, conditional on the fixation of a large-effect mutation at the locus.

**Figure S10:**
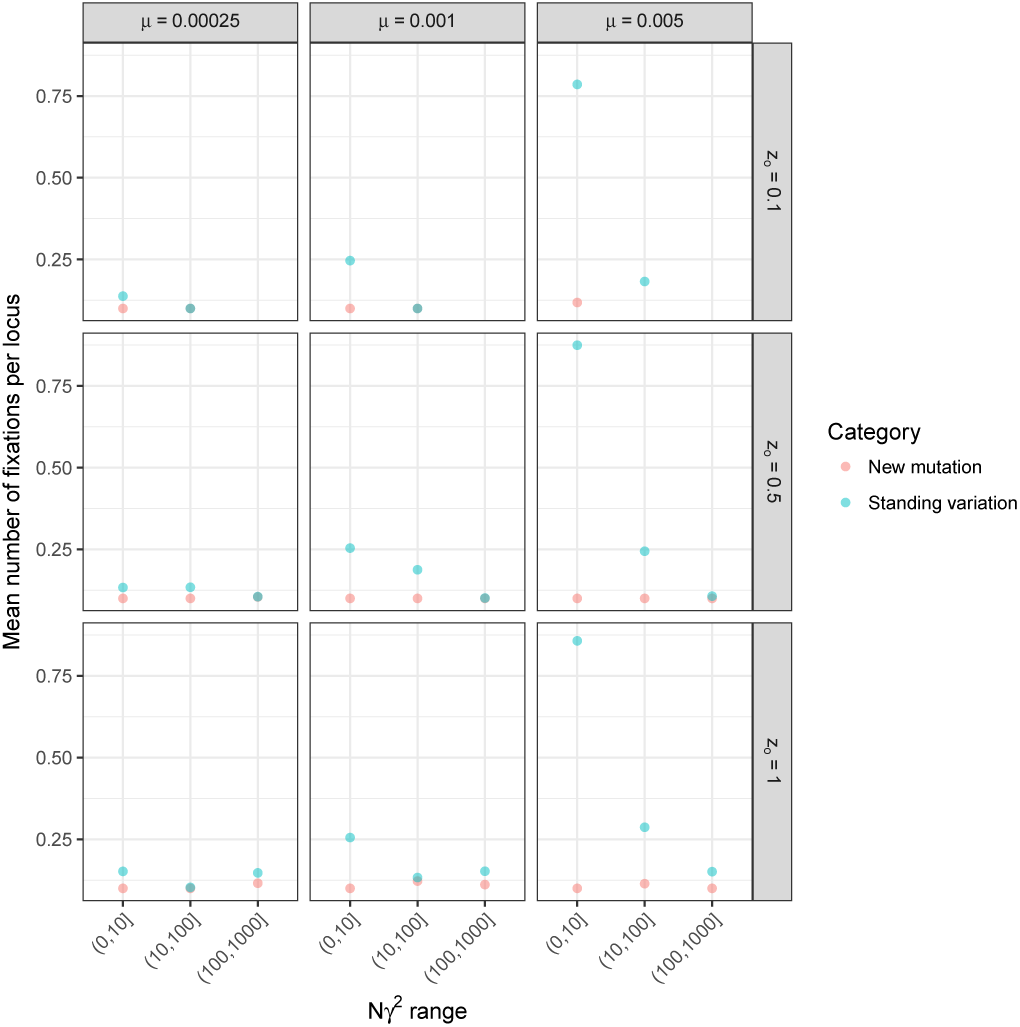
The mean number of fixations per locus. The means are taken for different ranges of *Nγ*^2^. Missing points mean no fixations in the relevant range.

**Figure S11:**
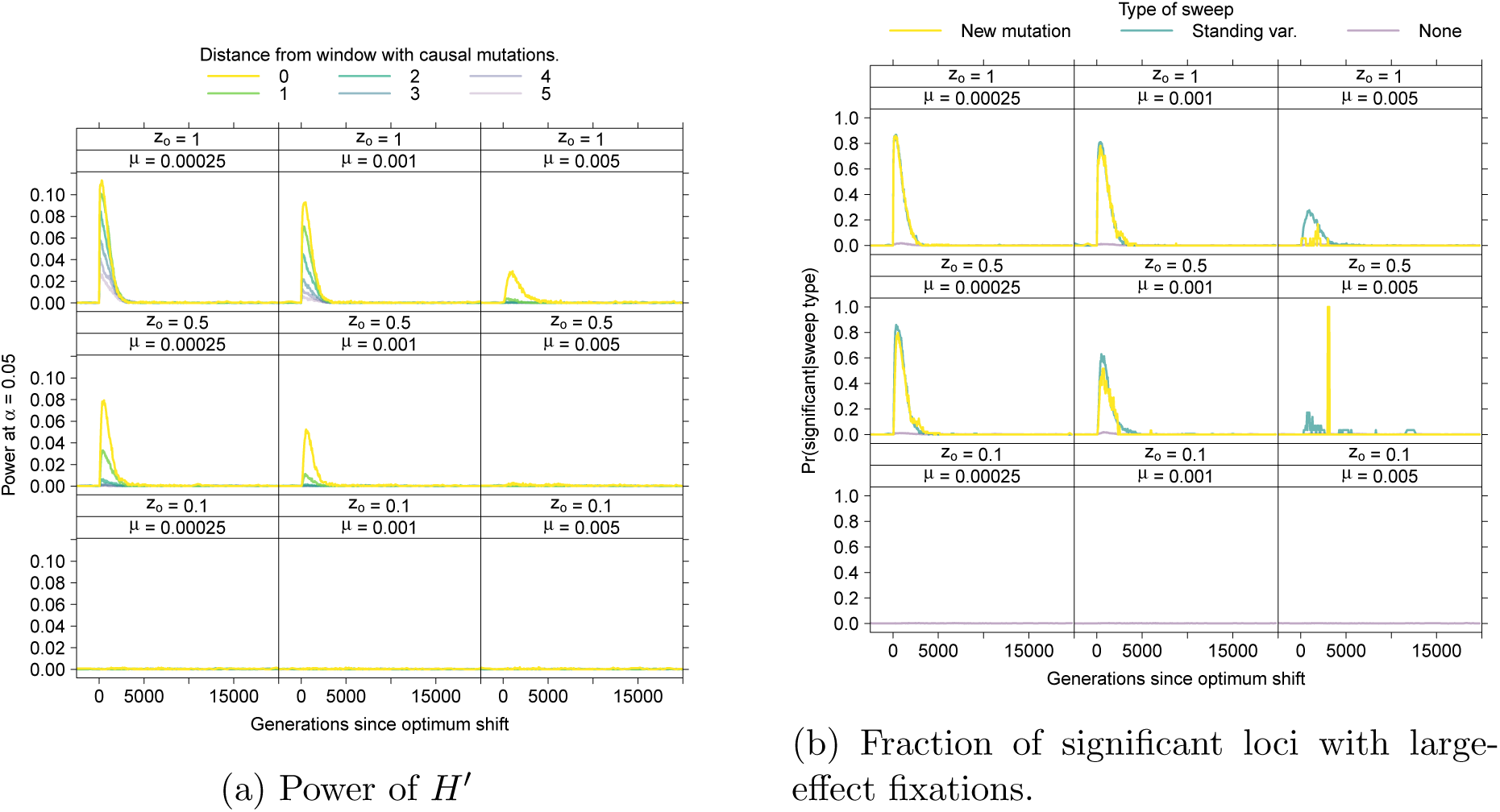
Power of *H′* (Zeng *et al.*, 2006) to reject the standard neutral model. S11a The power to reject the null model in windows of increasing distance from selected site as a function of time since the optimum shift, corrected so that the per-window rejection rate under the null model is 0.05. The windows are labeled using the same scheme as Figure S8. S11b The probability of a “significant” window at a locus, conditional on the fixation of a large-effect mutation at the locus.

**Figure S12:**
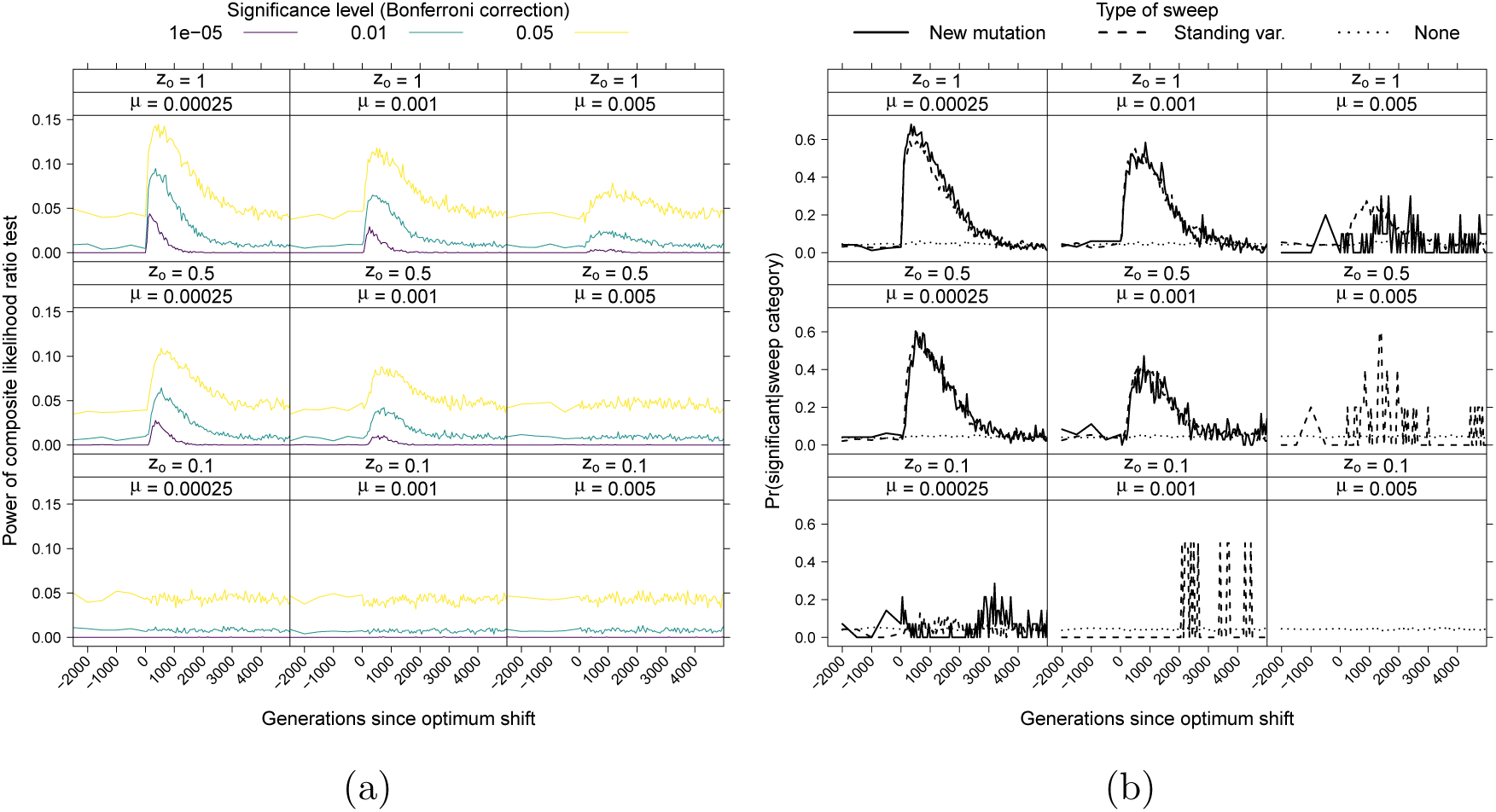
Power of the composite likelihood ratio test (CLRT) to reject the null model. All results are based on 10^6^ coalescent simulations using `msprime` (Kelleher *et al.*, 2016) and analyzed using SweeD (Pavlidis *et al.*, 2013). S12a The power to reject the null model at three different significance levels, after application of a Bonferonni correction. S12b The probability of a significant CLRT, conditional on the type of sweep.

**Figure S13:**
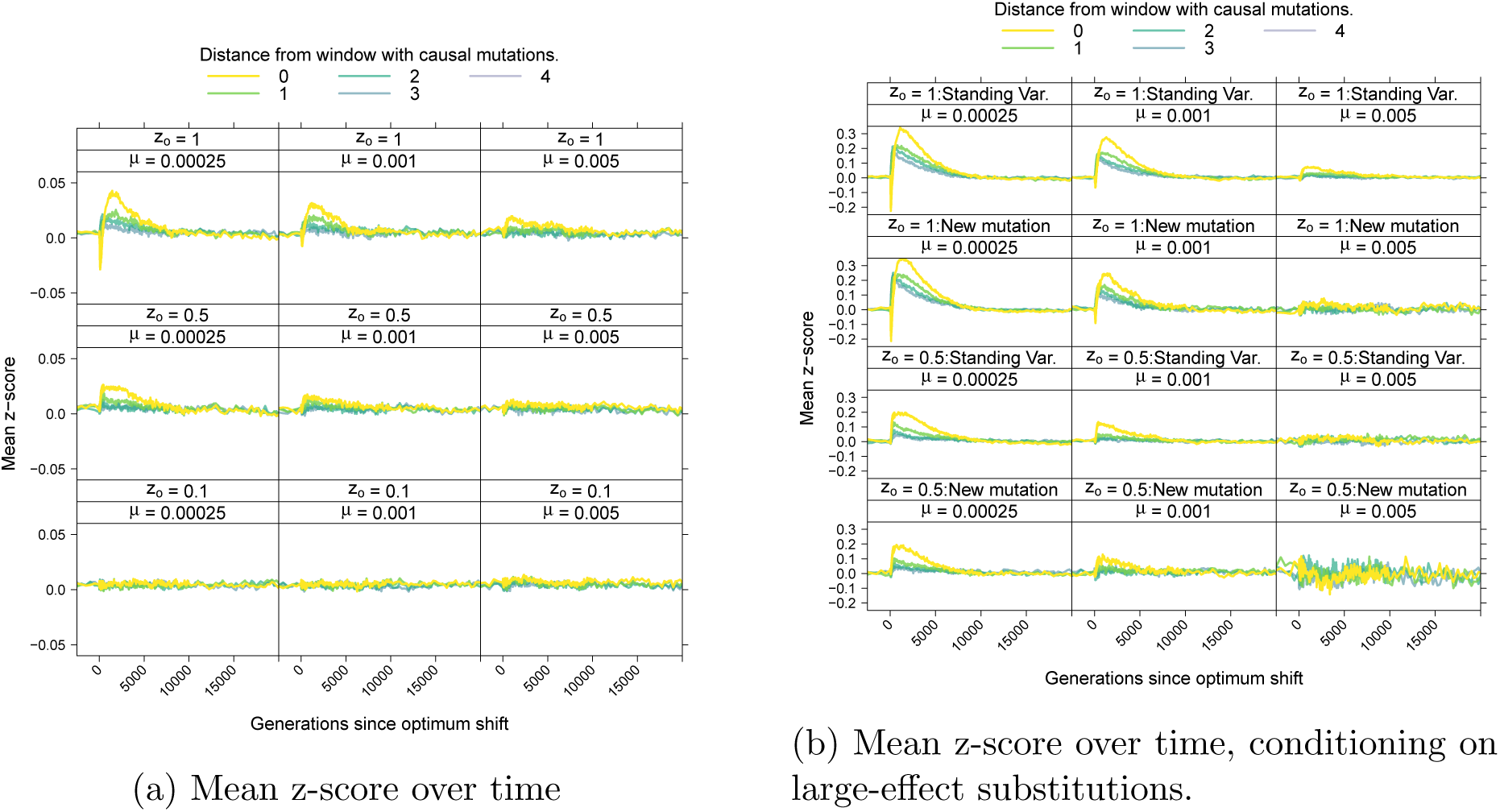
The mean z-score corresponding to the *nS*_*L*_ statistic (Ferrer-Admetlla *et al.*, 2014) over time. The data are plotted for different windows according to the scheme shown in Figure 2. S13a shows the average value over time and S13b shows the average value conditioning on fixation of large effect from standing variation or from a new mutation.

**Figure S14:**
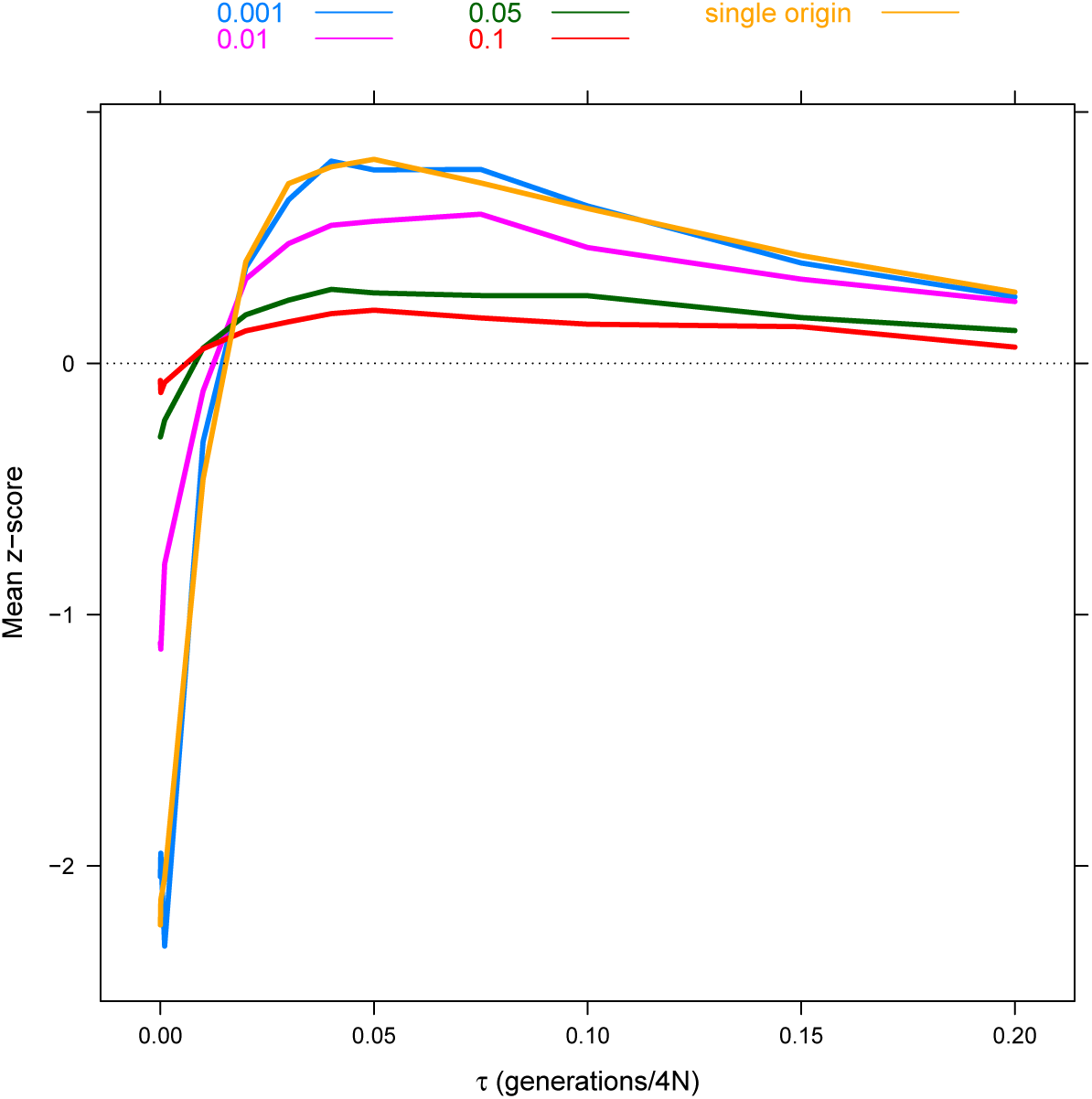
The average z-score associated with the *nS*_*L*_ statistic (Ferrer-Admetlla *et al.*, 2014) as a function of the time since the fixation of a strongly selected mutation. All lines are based on 100 replicates of a coalescent simulation (Kern and Schrider, 2016) of a sample size of 100 haplotypes and a region of size *θ* = *ρ* = 100. A single sweep with strength of selection *α* = 2*Ns* occurs in the middle of the region. The time since the fixation (*τ*) and the initial starting frequency of the mutation vary. A sweep from a single origin is a “hard sweep” with initial frequency 1*/*(2*N*_*e*_).

**Figure S15:**
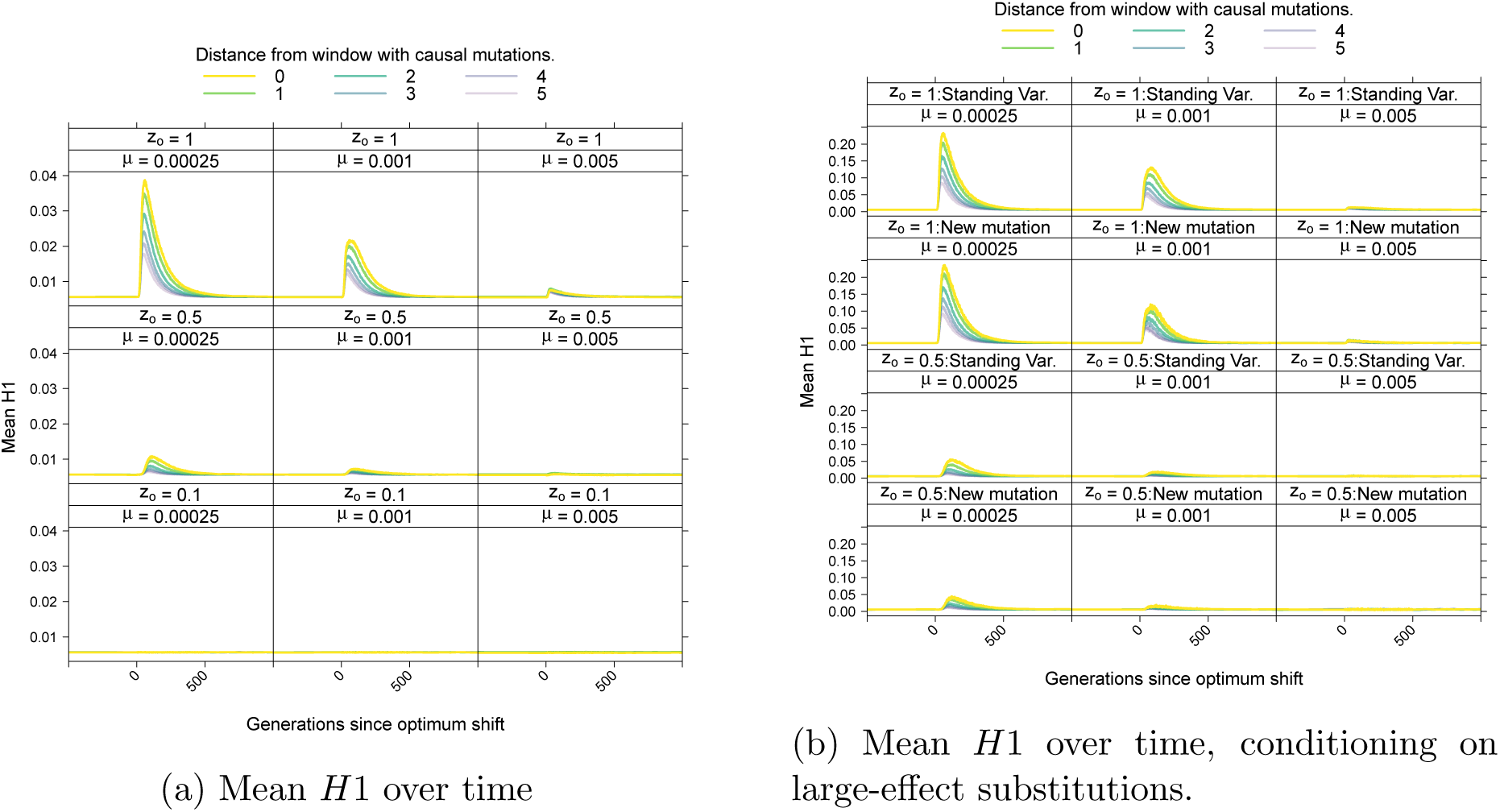
The mean *H*1 statistic (Garud *et al.*, 2015) over time. S15a shows the average value over time and S15b shows the average value conditioning on a fixation of large effect either from standing variation or from a new mutation.

**Figure S16:**
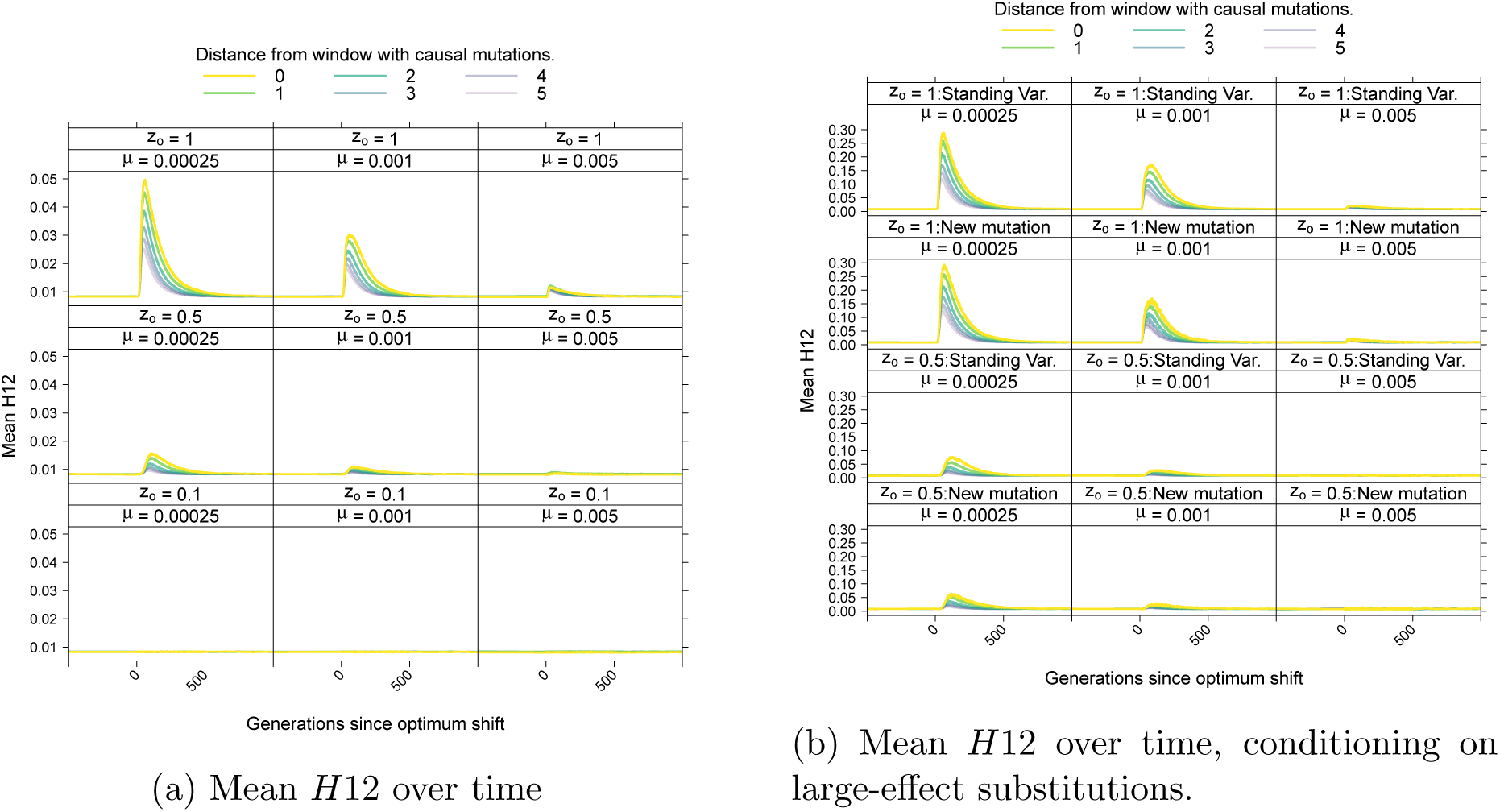
The mean *H*12 statistic (Garud *et al.*, 2015) over time. S16a shows the average value over time and S16b shows the average value conditioning on a fixation of large effect either from standing variation or from a new mutation.

**Figure S17:**
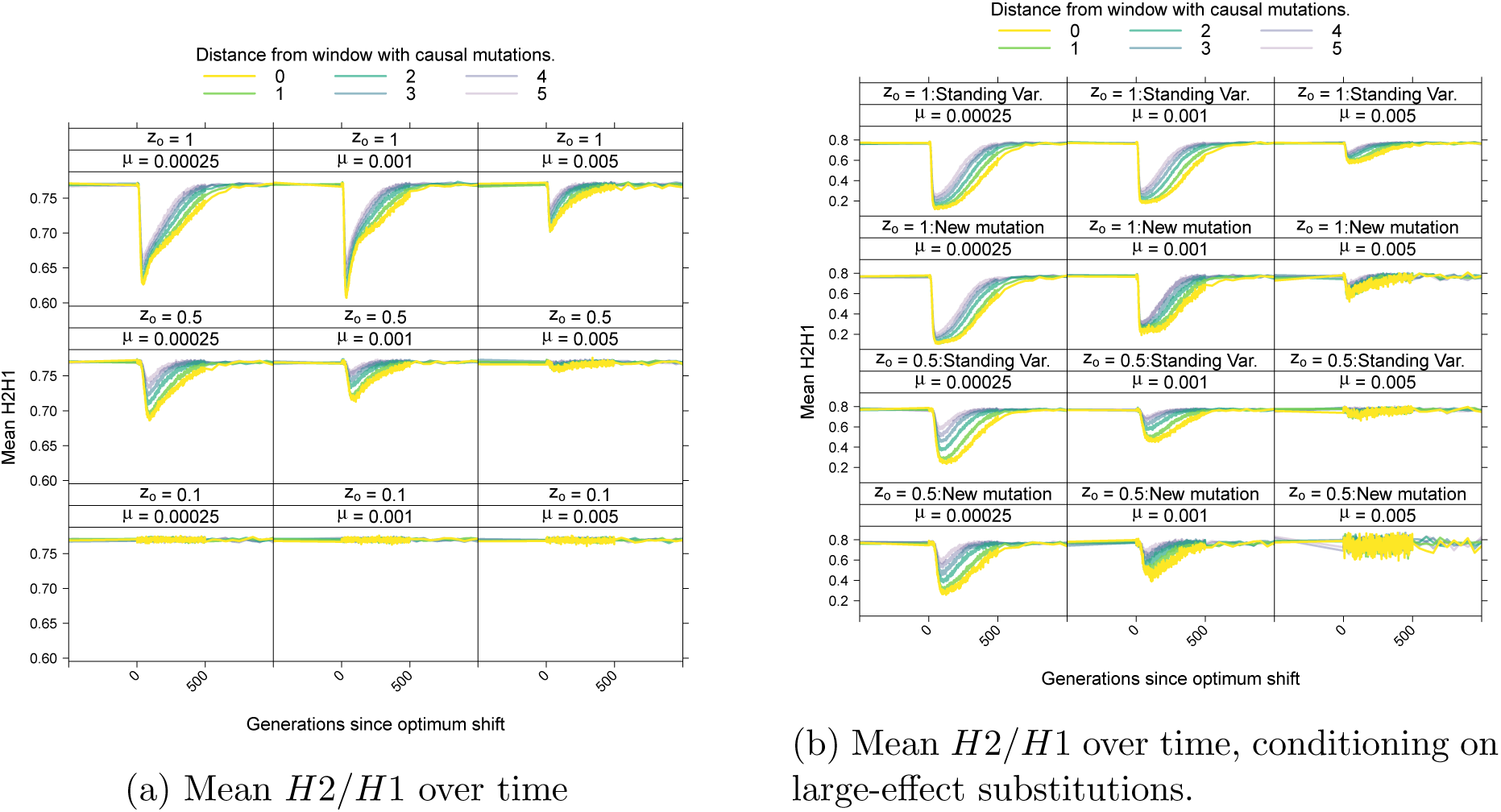
The mean *H*2*/H*1 statistic (Garud *et al.*, 2015) over time. S17a shows the average value over time and S17b shows the average value conditioning on a fixation of large effect either from standing variation or from a new mutation.

**Figure S18:**
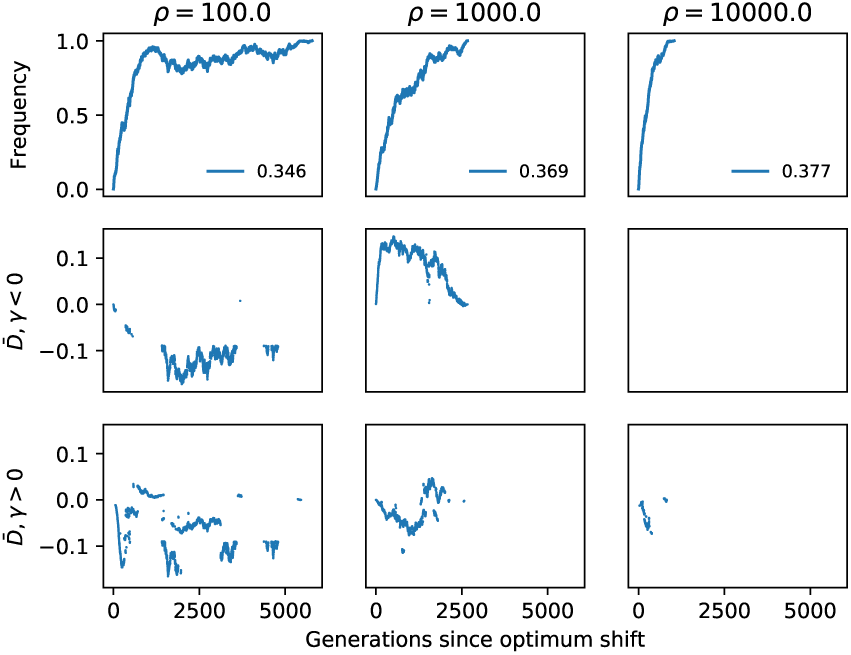
Within-locus linkage disequilibrium with fixations. Each column represents a single simulation replicate of a ten-locus system with *µ* = 5 × 10^−3^ and a different value of *ρ* = 4*Nr* per locus (column labels). Here, each simulation happens to have one fixation from a new mutation, whose frequency trajectory is shown in the top row and the effect sizes are shown in the panel legends. The second and third rows show the mean value of *D* = *p*_*AB*_ *– p*_*A*_*p*_*B*_, where *p*_*A*_ is the frequency of the sweeping variant and *p*_*B*_ is the frequency of another variant affecting trait values at the same locus. *D* is calculated after filtering variants with minor allele frequencies less than 0.1. The middle row shows 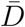 with respect to loci with effect sizes *γ >* 0 and the third row is with respect to *γ >* 0. Gaps in the 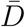 data mean that no mutations passing the frequency filter were present in that generation.

**Figure S19:**
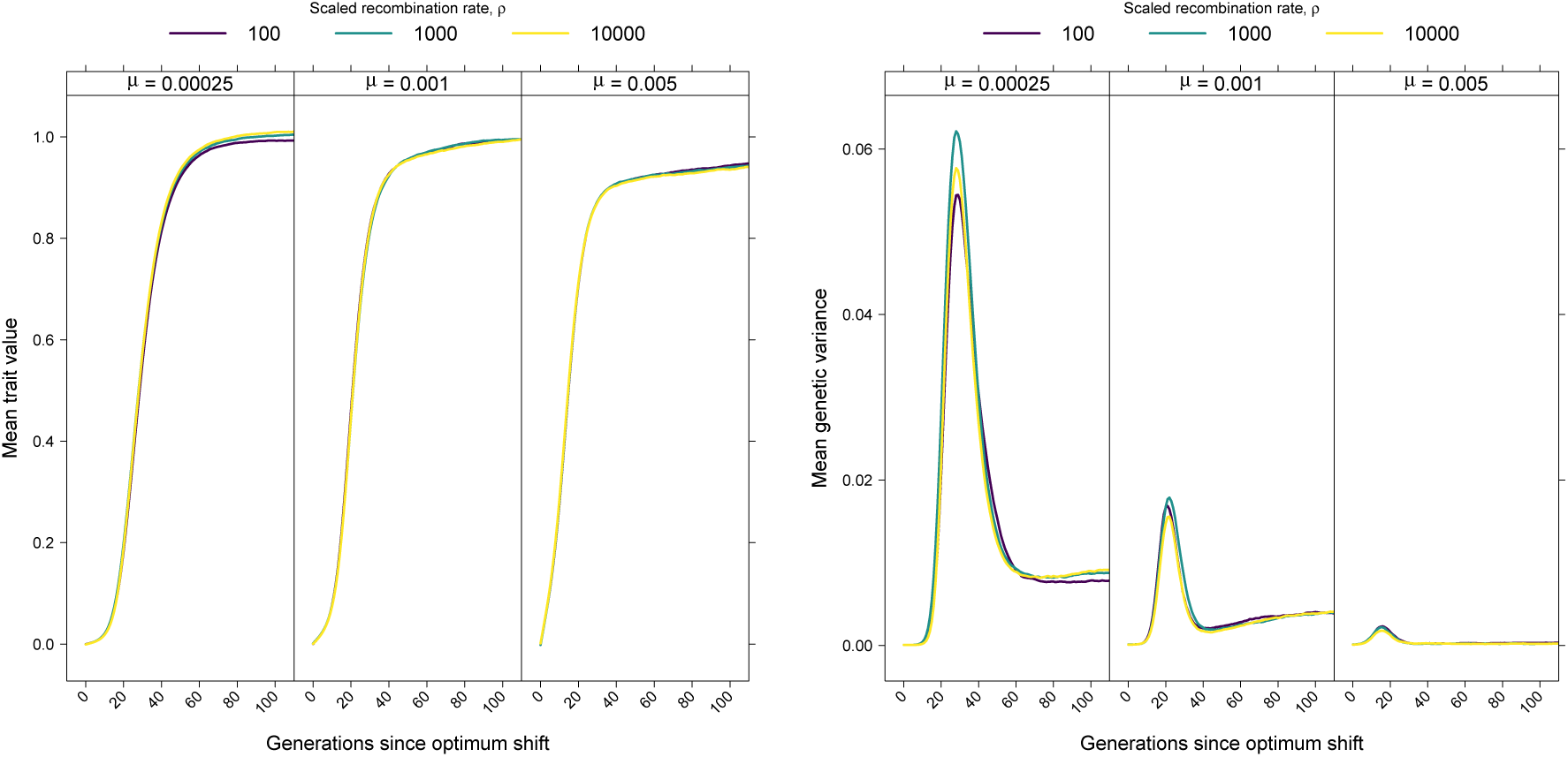
The mean genetic value and variance over time in a ten-locus system as the within-locus recombination rate *ρ* = 4*Nr* varies.

**Figure S20:**
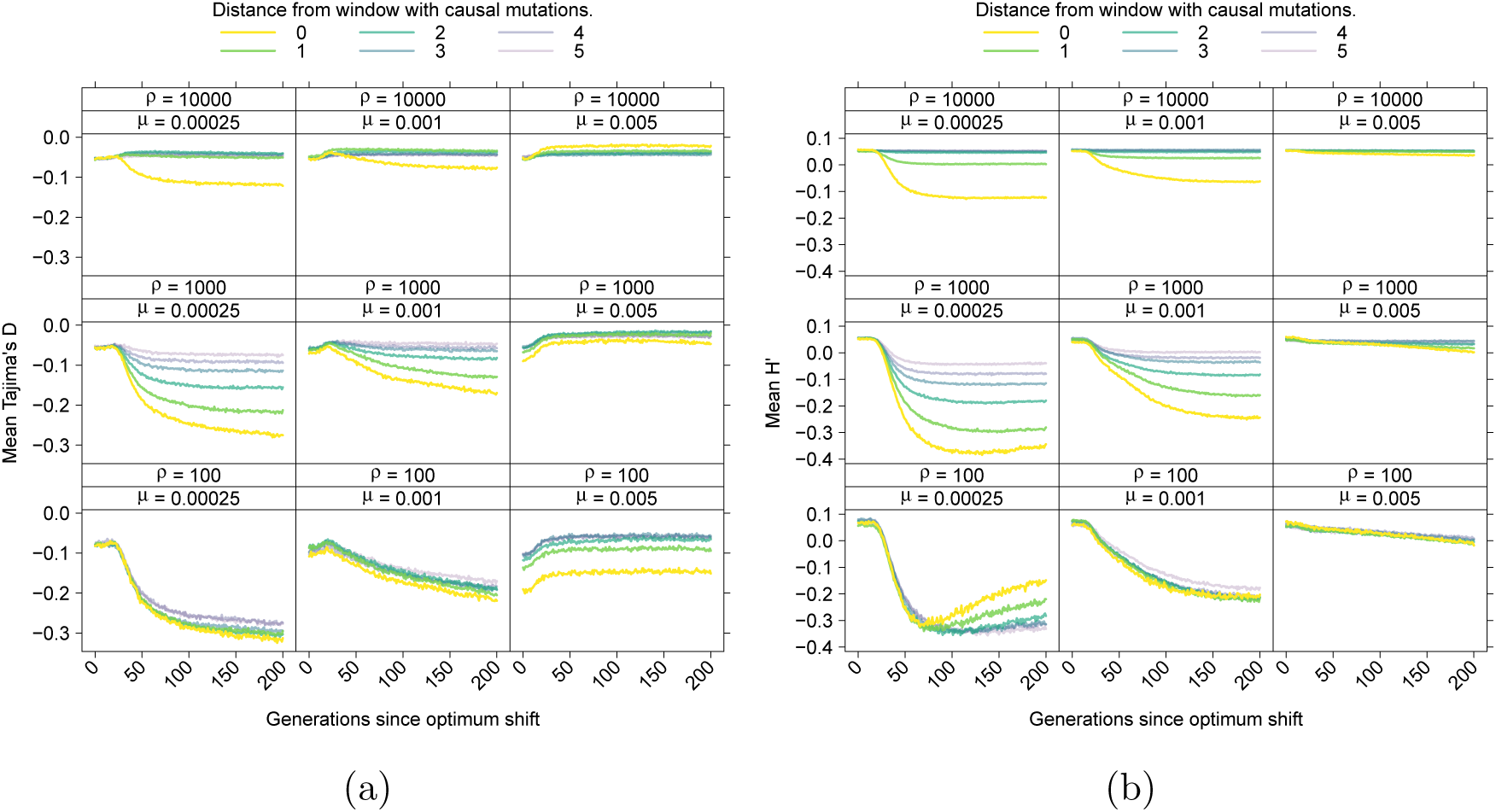
The mean behavior of Tajima’s (1989) *D* (a) and *H′* (Zeng *et al.*, 2006) (b) over time as the recombination rate within a locus, *ρ* = 4*Nr*, is varied. The layout of each panel is as in Figure 8, averaging across loci with the same distance from the central window of each locus (Figure 2) each generation.

**Figure S21:**
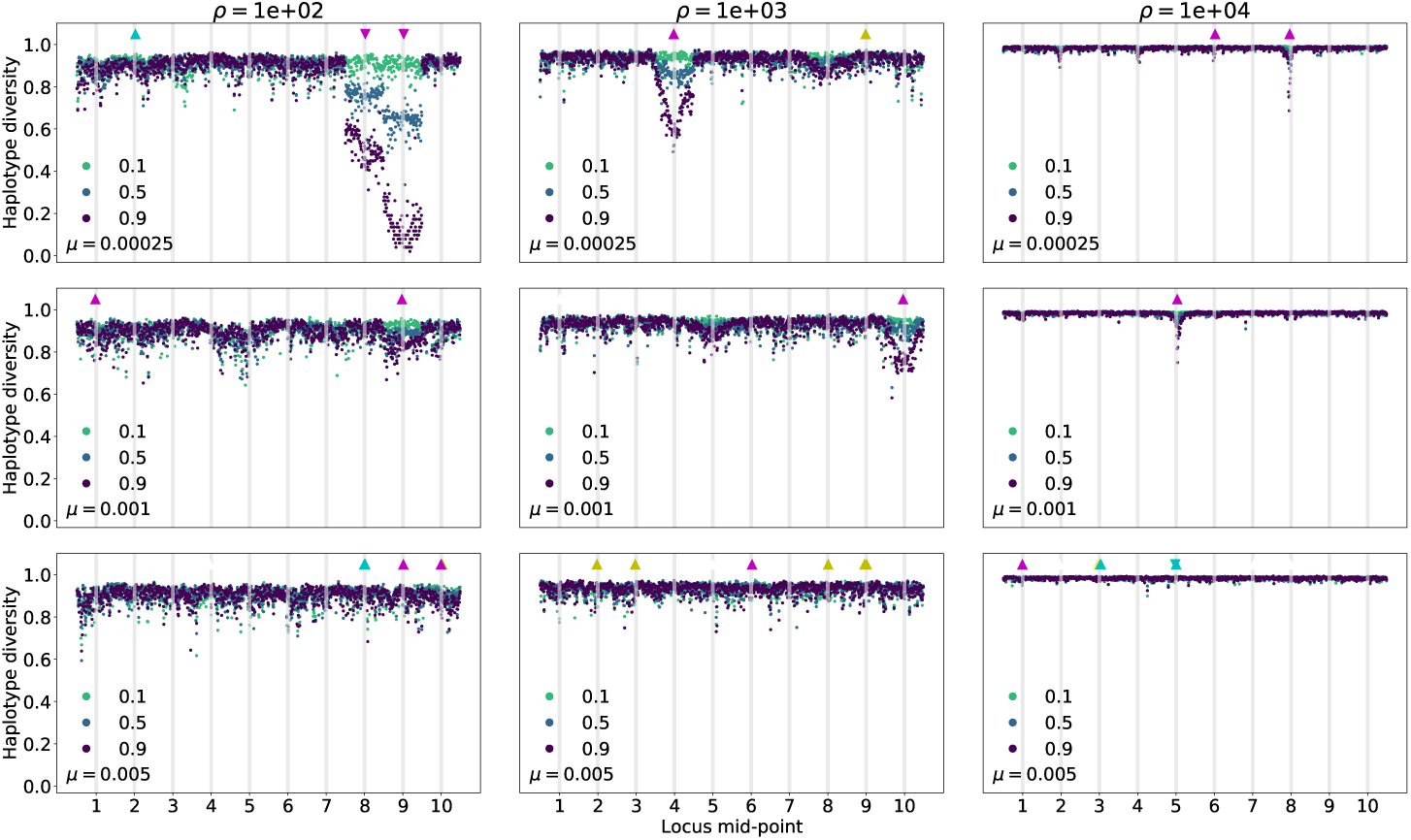
Haplotype diversity along the genome in a ten locus system as the within-locus recombination rate varies. Haplotype diversity is shown for three replicate simulations adapting to an optimum trait value of *z*_*o*_ = 1. The center column shows the same data as the final row of Figure 7. The left and right column decrease and increase *ρ* to 100 and 10,000, respectively. The triangles along the top of each panel show where fixations occurred. Triangles pointing up are fixations from standing variation, and those pointing downwards are fixations from new mutations arising within 100 generations of the optimum shift. Magenta refers to fixations with scaled effect sizes *Nγ*^2^ *>* 100, cyan indicates 10 *< Nγ*^2^ *≤* 100, and yellow refers to 1 *≤ Nγ*^2^ *≤* 10.

**Figure S22:**
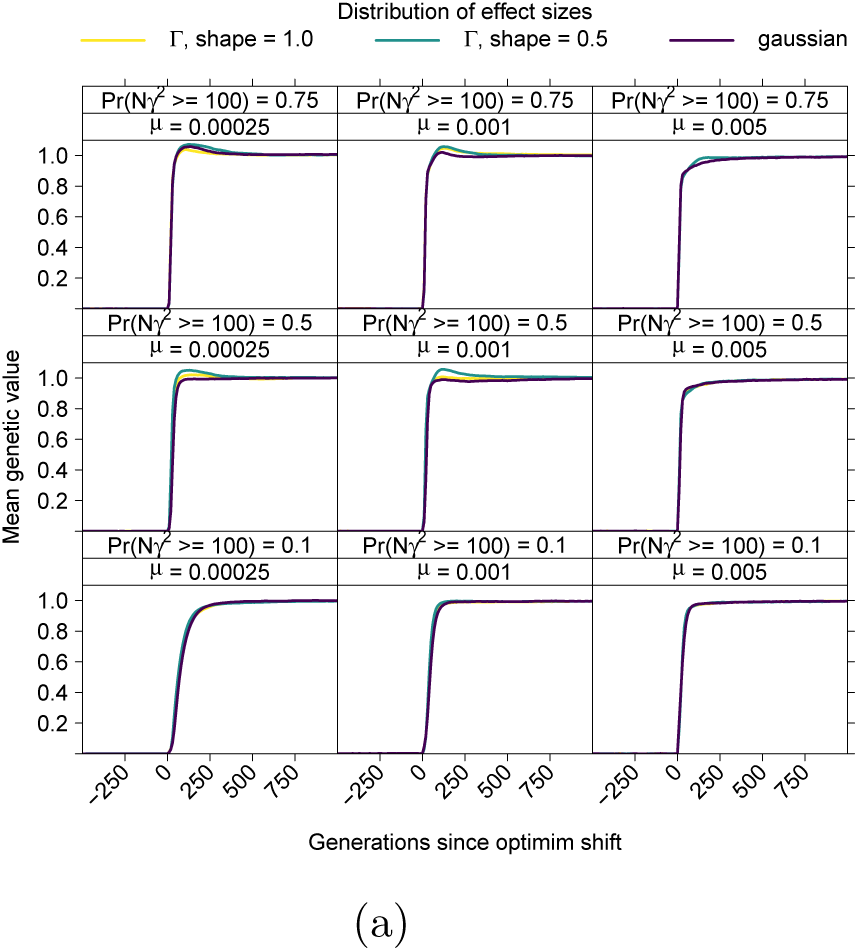
The average dynamics of adaptation as a function of the fraction of newly arising mutations of large effect. For a given mutation rate, *µ*, the probability that a new mutation has an effect size *Nγ*^2^ ≥ 100 is varied for three different distributions of effect sizes (DES), the Gaussian distribution and Gamma distributions with two difference shape parameters. S22a The mean trait value over time.

**Figure S23:**
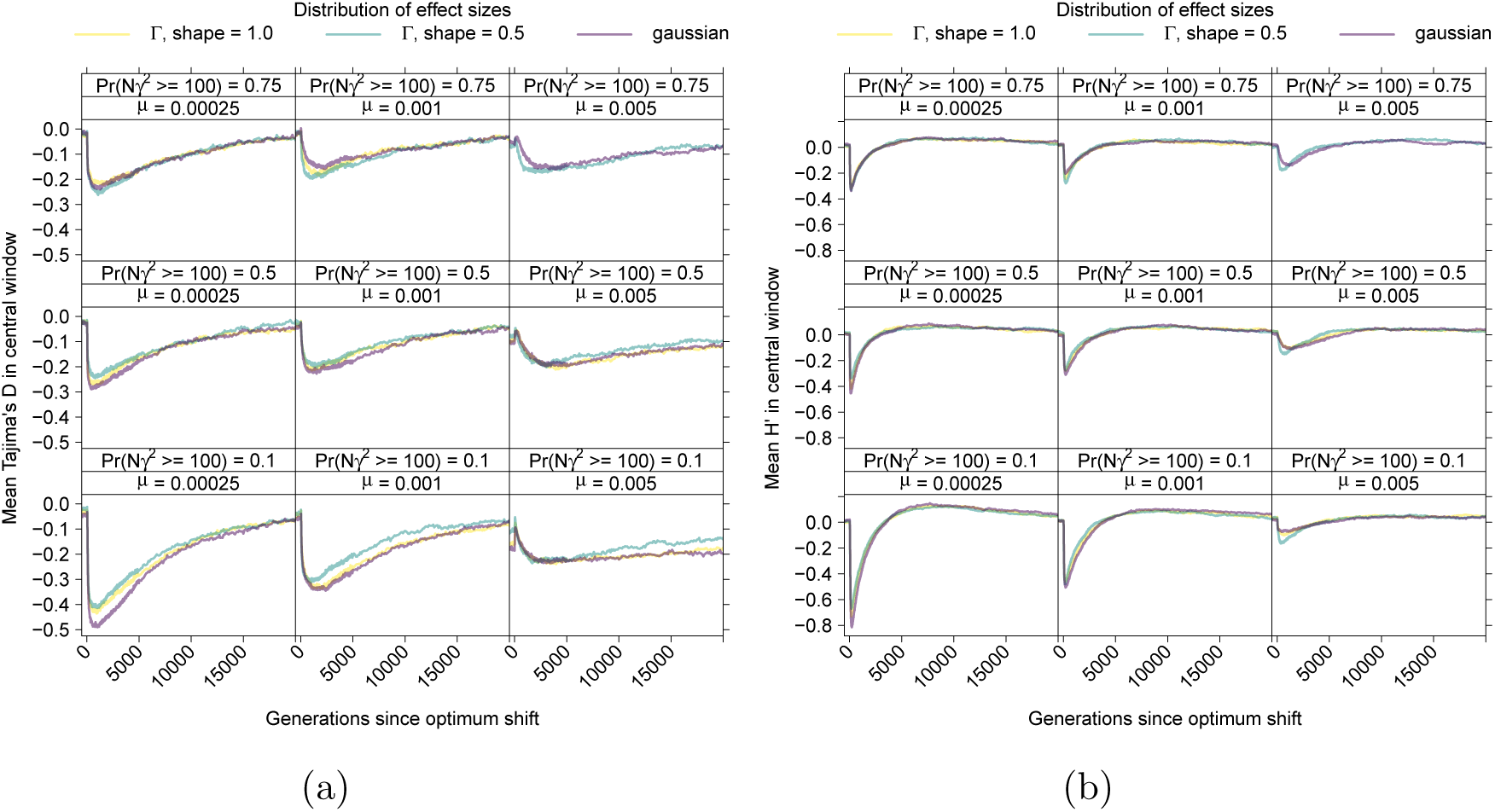
The average deviation from equilibrium patterns of variation as a function of the probability that a new mutation is of large effect. As in Figure S22, the mutation rate is held constant for three different distributions as *Pr*(*Nγ*^2^ ≥ 100) varies. S23a The mean of Tajima’s (1989) *D* over time in the central window of a locus, which is where mutations affecting the trait arise (Figure 2). S23b The mean *H′* (Zeng *et al.*, 2006) over time in the central window of a locus.

**Figure S24:**
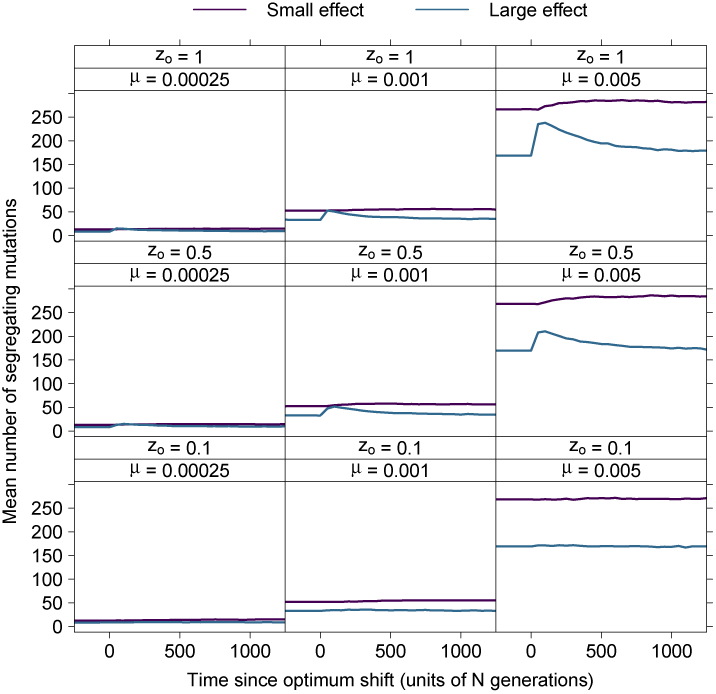
The mean number of segregating mutations affecting the trait over time in ten locus simulations. The different lines are for mutations of small effect (*Nγ*^2^ *<* 100) and large-effect variants (*Nγ*^2^ ≥ 100).

